# Combinatory EHMT and PARP inhibition induces an interferon response and a CD8 T cell-dependent tumor regression in PARP inhibitor-resistant models

**DOI:** 10.1101/2023.02.23.529773

**Authors:** Lily L. Nguyen, Zachary L. Watson, Raquel Ortega, Elizabeth R. Woodruff, Kimberly R. Jordan, Ritsuko Iwanaga, Tomomi M. Yamamoto, Courtney A. Bailey, Abigail D. Jeong, Saketh R. Guntupalli, Kian Behbakht, Veronica Gbaja, Nausica Arnoult, Edward B. Chuong, Benjamin G. Bitler

**Affiliations:** Molecular Cellular Developmental Biology, The University of Colorado Boulder, Boulder, CO 80309, USA; Department of Immunology and Microbiology, The University of Colorado Anschutz Medical Campus, Aurora, CO 80045, USA; Department of Obstetrics & Gynecology, Division of Gynecologic Oncology, The University of Colorado Anschutz Medical Campus, Aurora, CO 80045, USA; Division of Reproductive Sciences, Department of Obstetrics and Gynecology, University of Colorado School of Medicine, Aurora, CO, 80045; Epizyme/Ipsen Company, Cambridge, MA

## Abstract

Euchromatic histone lysine methyltransferases 1 and 2 (EHMT1/2), which catalyze demethylation of histone H3 lysine 9 (H3K9me2), contribute to tumorigenesis and therapy resistance through unknown mechanisms of action. In ovarian cancer, EHMT1/2 and H3K9me2 are directly linked to acquired resistance to poly-ADP-ribose polymerase (PARP) inhibitors and are correlated with poor clinical outcomes. Using a combination of experimental and bioinformatic analyses in several PARP inhibitor resistant ovarian cancer models, we demonstrate that combinatory inhibition of EHMT and PARP is effective in treating PARP inhibitor resistant ovarian cancers. Our *in vitro* studies show that combinatory therapy reactivates transposable elements, increases immunostimulatory dsRNA formation, and elicits several immune signaling pathways. Our *in vivo* studies show that both single inhibition of EHMT and combinatory inhibition of EHMT and PARP reduces tumor burden, and that this reduction is dependent on CD8 T cells. Together, our results uncover a direct mechanism by which EHMT inhibition helps to overcome PARP inhibitor resistance and shows how an epigenetic therapy can be used to enhance anti-tumor immunity and address therapy resistance.

## INTRODUCTION

Ovarian cancer is the deadliest gynecologic malignancy and has a high propensity to develop resistance to chemo- and targeted therapies^1^. Poly-ADP-ribose-polymerase (PARP) inhibitors, a targeted therapy, are used as first-line maintenance therapies for ovarian cancer due to their efficacy against cells with homologous recombination (HR) deficiencies (e.g., BRCA1/2-mutations)^2^. PARP inhibitors inhibit single-stranded DNA repair, resulting in double-stranded DNA (dsDNA) breaks that cannot be repaired in HR deficient cells, therefore inducing DNA-damage-induced apoptosis^3, 4^.

Recently, PARP inhibitors have also been shown to be clinically beneficial in HR proficient tumors^5^. The damaged DNA induced by PARP inhibition results in dsDNA formations that can trigger the cGAS-STING pathway^6, 7^. This pathway stimulates type I interferon signaling and subsequently activates anti-tumor immunity^6, 7^. Additionally, PARP inhibitors are now FDA-approved for breast, prostate, and pancreatic cancers^8–10^. However, resistance to PARP inhibitors occurs in approximately 50% of patients and there is an unmet need to develop therapeutic strategies that can combat PARP inhibitor resistance^11, 12^.

Epigenetic therapies have been shown to resensitize some cancer cells to chemo- or targeted-therapies^13^. DNA methyltransferase (DNMT) inhibitors such as decitabine and 5-Azacitidine can resensitize ovarian cancer to platinum therapy^14^. Histone deacetylase (HDAC) inhibitors resensitize sorafenib-resistant hepatocellular carcinoma to therapy^15^ and many other combinations with epigenetic therapies are currently in clinical trials^16^. However, many of these epigenetic therapies have global, non-specific targets and can have devastating side effects with minimal clinical improvements^17^. Due to the potential benefits of epigenome reprogramming, including the reversibility of the chromatin modifications, it is important to explore other epigenetic therapies that may have a more specific mechanism of action. To that end, we have shown that inhibitors of euchromatic histone lysine methyltransferases 1 and 2 (EHMT1/2) can resensitize ovarian cancer cells to PARP inhibitor treatment^18^. EHMT1 and EHMT2 are chromatin “readers” and “writers” that catalyze mono- and di- methylation of histone H3 lysine 9 (H3K9) residues^19^. Overexpression of EHMT1 and EHMT2 correlates with poorer survival outcomes, increased tumorigenesis, and therapy resistance in several cancer types^20^. In PARP inhibitor resistant ovarian cancer cells, both EHMT1/2 and H3K9me2 correlate with poorer survival outcomes and are causally linked to PARP inhibitor resistance^18^.

Though there are suggestions on how overexpression of EHMT1 and/or EHMT2 confers poorer outcomes (e.g. increased epithelial mesenchymal transition, maintenance of stemness, and augmented DNA repair), it is appreciated that these results are likely context-dependent^20^.

In addition to gene expression regulation, epigenetic inhibitors also reactivate transposable elements (TEs)^21^. Transposable elements are repetitive DNA segments that are normally repressed through various epigenetic mechanisms including H3K9me2^22, 23^. When these epigenetic modifications are removed, such as through epigenetic inhibitors, TEs are de-repressed via loss of H3K9me2, transcribed, and form immunostimulatory double-stranded RNA (dsRNA)^24–27^. These host-derived dsRNA mimic viral dsRNA and are detected by intracellular sensors such as RIGI and MDA5, triggering an interferon response^24–26^. The interferon response in conjunction with an intact immune system can promote anti-tumor immunity through the recruitment and activation of T cells. This process, termed viral mimicry, has been studied in the context of DNMT inhibitors and several epithelial cancers^24^. However, viral mimicry has not been studied in the context of PARP inhibitor resistance and targeting EHMT1/2 activity.

Here, we investigated the mechanism of action of combinatory EHMT and PARP inhibition in *in vitro* and *in vivo* models of PARP inhibitor resistant ovarian cancer cells, including isogeneic cell lines, syngeneic immunocompetent mouse models, publicly available datasets, and primary ovarian cancer tumors. Our studies show that combinatory EHMT and PARP inhibition is effective in treating PARP inhibitor resistant ovarian cancer cells *in vitro* and *in vivo*. Responses to combinatory EHMT and PARP inhibition is partially dependent on RIGI and MDA5 *in vitro*, on enhancement of cytotoxic T cell activation *ex vivo*, and on CD8 T cell activity *in vivo*. Our data suggest that CD8 T cells may be activated by increased interferon response when EHMT activity and H3K9me2 were reduced through pharmacologic inhibition. Reactivation of transposable elements may be one source of increased immunostimulatory dsRNA triggering an interferon signaling cascade. The increase in interferon signaling may then recruits CD8 T cells and promotes anti-tumor immunity.

## RESULTS

### Combinatory EHMT and PARP inhibition induces transcriptional reprogramming and upregulation of interferon pathways

Compared to isogeneic PARP inhibitor (PARPi) sensitive cells, Watson *et al*^18^ has shown that PARPi resistance in ovarian cancer is linked to overexpression of EHMT1/2 and increased H3K9me2. Critically, an EHMT inhibitor, UNC0642, in combination with a PARP inhibitor, Olaparib, was found to be effective in treating PARPi-resistant cells *in vitro*. To determine the mechanism of action of the combination therapy, we measured the transcriptional changes induced by combinatory EHMT and PARP inhibition. We used RNA-seq to profile several PARPi-resistant ovarian cancer cell lines (PEO1-R, Kuramochi-R (Kura-R), OVCAR420-R) and therapy-naïve, patient-derived, primary tumors (*ex vivo* cultures^28^) treated with control (DMSO), a PARP inhibitor (2*μ*M Olaparib), an EHMT inhibitor (1*μ*M UNC0642), or combination (2*μ*M Olaparib and 1*μ*M UNC0642) therapy for 72 hours (Fig. 1A), as this is the minimal time needed to decrease H3K9me2 levels and before cell defects were observed (Fig S1A-C). As expected, when a repressive epigenetic mark is reduced, differential gene expression analysis^29^ showed that cells treated with single EHMT or combinatory EHMT and PARP inhibition upregulated expression of more genes than cells treated with DMSO or PARP inhibition alone (Fig. 1B-D, Fig. S1D-L). Gene set enrichment analysis (GSEA)^30, 31^ showed several interferon and immune related signaling pathways significantly enriched in combination treated cells compared to control or singly treated cells (Fig. 1E-G, Fig. S1M-R). This upregulation of interferon and immune signaling pathways was only observed in the BRCA2-mutated cell lines (PEO1-R and Kura-R), but not in the BRCA wildtype cells (OVCAR420) (Fig. 1E-G, Fig. S1M-R). We next focused on the interferon-*α* and interferon-*γ* pathways since they were consistently the most enriched in combination-treated cells (Fig. 1H-J). Intersecting individual genes upregulated in these two pathways, 79 genes were shared between the two BRCA2-mutated, PARPi-resistant cell lines and the patient-derived *ex vivo* tissues. (Fig. 1K). Among these genes includes CCL5, CXCL10, and CXCL11 (Fig. 1L). The mRNA expression of all three of these cytokines in patient tumors are correlated with improved overall survival (Fig. 1M), suggesting a beneficial role for interferon signaling in patient survival. Collectively, our RNA-seq analysis shows that combinatory EHMT and PARP inhibition upregulates interferon signaling pathways in both PARPi-resistant ovarian cancer cells and therapy-naive primary tumors.

**Figure 1:**
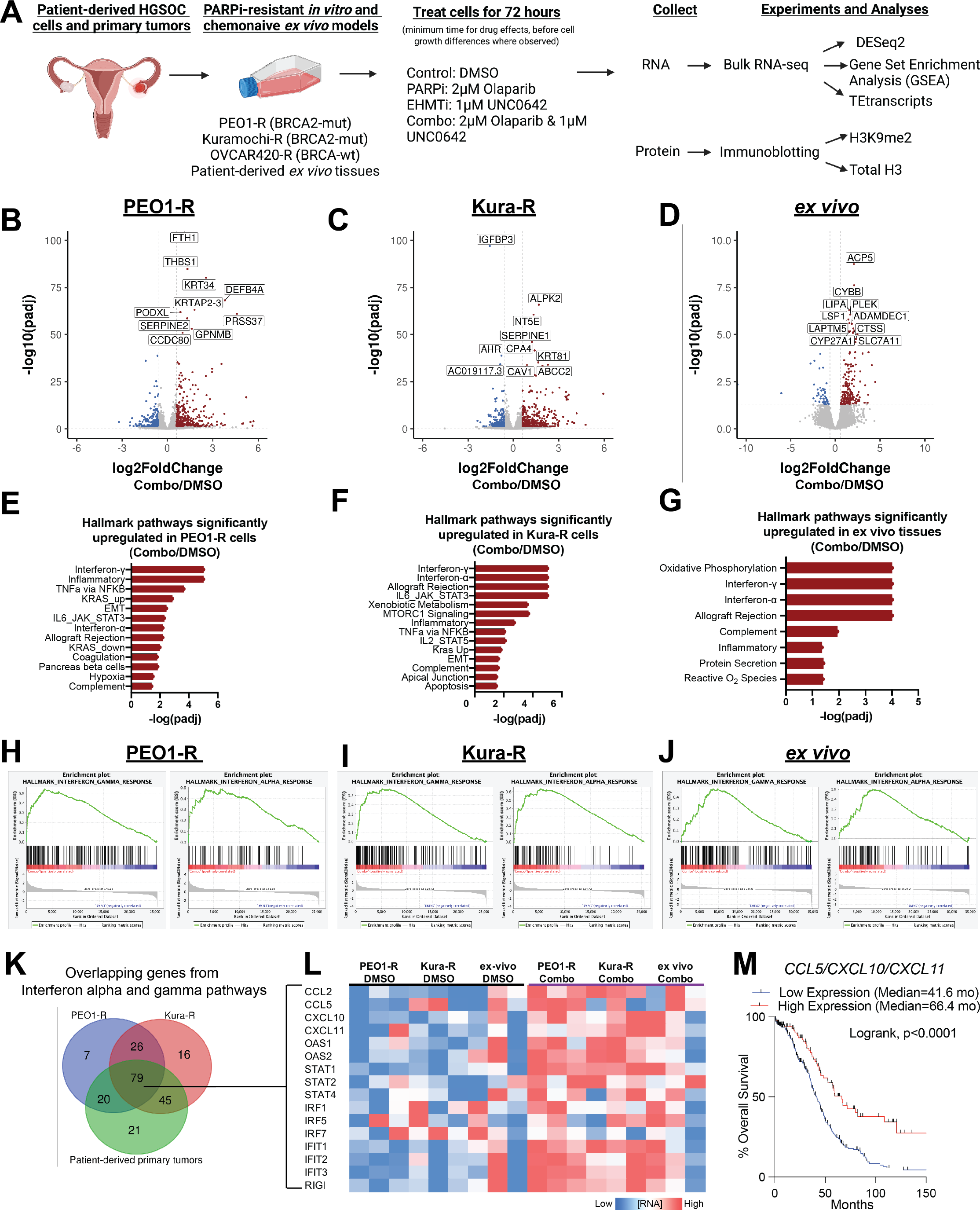
Combinatory EHMT and PARP inhibition induces transcriptional reprogramming and upregulation of interferon pathways. **A)** Schematic of experimental details, sequencing, and analysis workflow **B-D)** Volcano plots of differentially regulated genes in PEO1-R (B), Kura-R (C), and patient-derived *ex vivo* tissue (D) comparing combination treatment (Olaparib/UNC0642) to control (DMSO). **E-F)** Hallmark pathways upregulated in PEO1-R (E), Kura-R (F), and patient-derived *ex-vivo* tissue (G) treated with Olaparib/UNC0642 compared to DMSO. **H-J)** Enrichment plots of interferon-*α* and interferon-*γ* pathways in PEO1-R (H), Kura-R (I), and patient-derived *ex-vivo* tissue (J) treated with Olaparib/UNC0642 compared to DMSO. **K)** Venn diagram of overlapping genes from interferon-*α* and -*γ* pathways. **L)** Heatmap of 16 out of the 79 overlapping genes from **K)**. **M)** Overall survival Kaplan-Meier of patients with ovarian cancer with high or low CCL5/CXCL10/CXCL11 mRNA expression. Based on the median expression of each. Statistical test, Log-rank. *p<0.05.

### Combination therapy induces expression of transposable elements

The dramatic increase in interferon signaling in several PARPi-resistant cell lines and patient-derived *ex vivo* samples after treatment with an epigenetic inhibitor is consistent with a growing body of evidence that many epigenetic inhibitors activate interferon signaling by transcriptionally reactivating transposable elements (TEs), which can form immunostimulatory dsRNAs^24–27^. Induced dsRNAs are detected in the cells by intracellular sensors such as RIGI and MDA5, which stimulates the interferon signaling cascade. While dsRNAs can be generated by many sources, TEs are a potent and well-established source of dsRNAs that are activated upon epigenetic inhibitor treatment^25, 26^. To determine if any of the treatments induce interferon signaling through upregulation of TEs, we used TEtranscripts^32^ to analyze changes in TE family-level transcription. In PEO1-R and Kura-R cell lines and in *ex vivo* cultures of primary tumors, we found that combinatory EHMT and PARP inhibition significantly induced transcription of multiple TE families compared to control (Fig. 2A-C) while single treatments did not significantly induce any TE families (Fig. S2A-C, E-F). In OVCAR420-R cells, both single EHMT and combinatory EHMT and PARP inhibition induced multiple TE families compared to control (Fig S2D, H-I). We observed induction of both endogenous retrovirus and Alu repeat families, including LTR8B, LTR18B, HERVL18, AluJ, and AluS.

**Figure 2:**
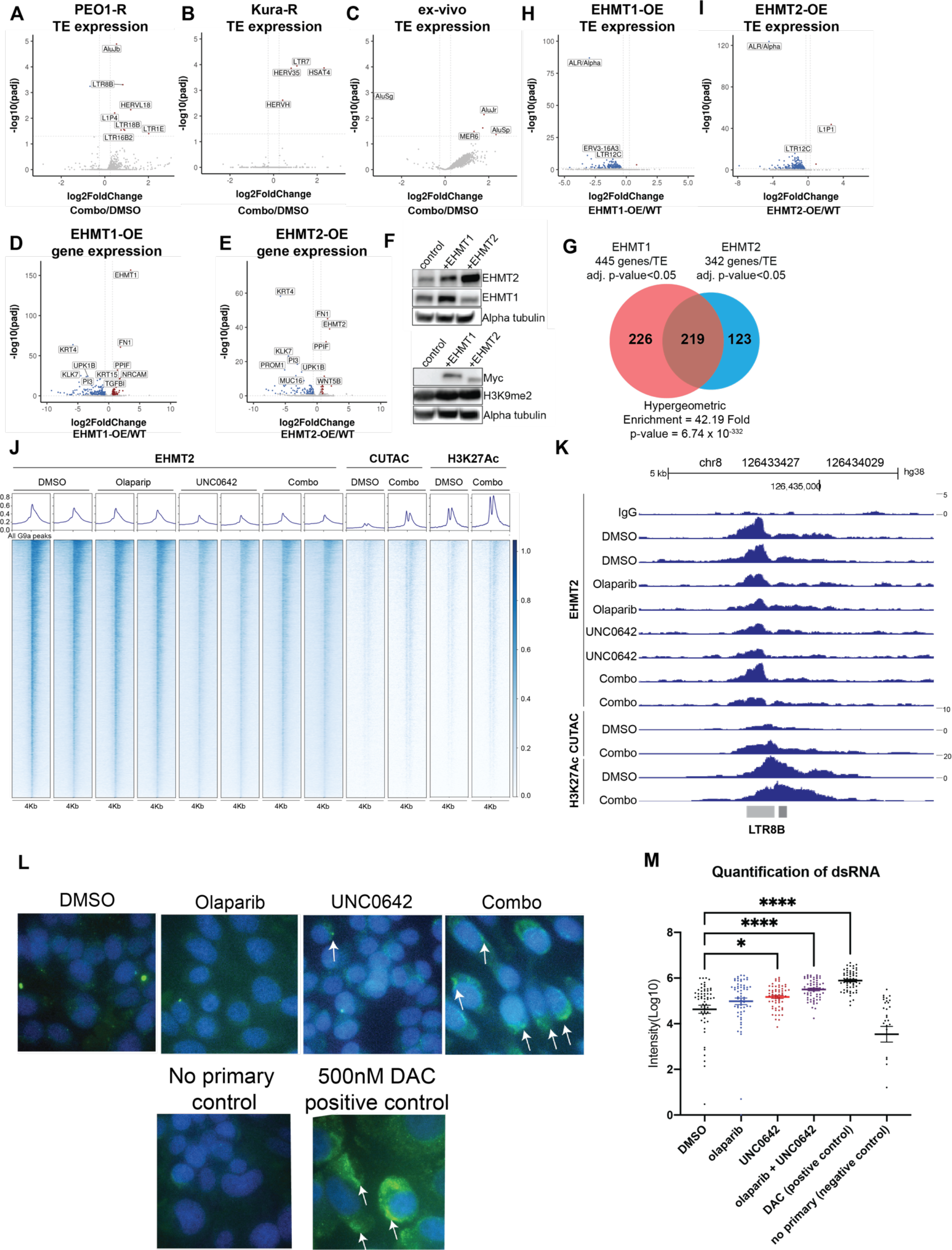
Combination therapy induces expression of transposable elements. **A-C)** Volcano plots of transposable element (TE) families with differentially expressed transcripts in PEO1-R (A), Kura-R (B), and patient-derived *ex vivo* tissues (C) treated with Olaparib/UNC0642 compared to DMSO control. **D-E)** Volcano plot of differentially regulated genes in EHMT1 (D) and EHMT2 (E) overexpressed PEO1 cell lines compared to wildtype PEO1 cells. **F)** Protein from EHMT1 and EHMT2 overexpressed PEO1 cells used for immunoblotting against EHMT1, EHMT2, Myc, and H3K9me2. Loading control, Alpha tubulin. **G)** Venn diagram of differentially regulated genes and TE in EHMT1 and EHMT2 overexpressed PEO1 cells. **H-I)** Volcano plot of TE families with differentially expressed transcripts in EHMT1 (H) AND EHMT2 (I) overexpressed PEO1 cells compared to wildtype PEO1 cells. **J)** Heatmap of EHMT2, CUTAC (open chromatin), and H3K27Ac (enhancer) signals in PEO1-R cells treated with DMSO, Olaparib, UNC0642, or Olaparib/UNC0642. **K)** Genome browser view of an LTR8B element. **L)** Representative immunofluorescence images of dsRNA in PEO1-R cells treated with DMSO, Olaparib, UNC0642, or Olaparib/UNC0642 for 72 hours. White arrows = positive signal. **M)** Quantification of immunofluorescence images from panel L (not all significant comparisons are shown). Error bars, SEM. Statistical test, multicomparison ANOVA. *p<0.05, ****p<0.0001.

TEs are normally repressed through epigenetic mechanisms such as H3K9 methylation^22, 23^ so the observation that TE transcript levels are increased with combinatory EHMT and PARP inhibition but not with single EHMT inhibition was interesting. Although H3K9me2 has been associated with TE repression in human^33^, it is not known whether H3K9me2 repressive marks around TE loci are deposited by EHMT or by another histone methyltransferases. To determine if EHMT1/2 regulates TE transcript expression, we overexpressed EHMT1 and EHMT2 in PARPi-sensitive PEO1 cells. Bulk RNA-sequencing and immunoblotting showed that EHMT1 and EHMT2 were both overexpressed at the RNA (Fig. 2D-E) and protein levels (Fig. 2F). As expected, given the increased levels of a repressive epigenetic modifier, we found that more genes were differentially down-regulated than there were upregulated (Fig. 2D-E), and 219 out of the 568 differentially regulated genes were overlapped between EHMT1 and EHMT2 overexpressed cell lines (Fig. 2G). In addition, transcription of multiple TE families was significantly repressed in both EHMT1 and EHMT2 overexpressed cell lines compared to EHMT1/2 wildtype PEO1 cells (Fig. 2H-I).

After establishing a link between EHMT activation and TE transcript expression, we next determined whether EHMT1/2 directly regulates TEs using CUT&RUN to identify where on the genome EHMT2 is bound in PEO1-R cells treated with control (DMSO), a PARP inhibitor (2*μ*M Olaparib), an EHMT inhibitor (1*μ*M UNC0642), or combination (2*μ*M Olaparib and 1*μ*M UNC0642) therapy for 72 hours. We also used CUT&TAG to examine areas of open chromatin and enhancer activity in PEO1-R cells treated with control (DMSO) or combination (2*μ*M Olaparib and 1*μ*M UNC0642) therapy for 72 hours. The CUT&RUN experiment showed that EHMT2 indeed binds to the genome, and, interestingly, DNA-bound EHMT2 decreases with single and combinatory EHMT inhibition, indicating that UNC0642 can inhibit both the methyltransferase and DNA-binding activity (Fig. 2J). Additionally, we observed increased open chromatin and H3K27Ac enhancer marks at EHMT2 sites when cells were treated with combination therapy (Fig. 2J). Next, we asked whether specific TE families were significantly enriched at EHMT2 sites by GIGGLE analysis^34, 35^. Indeed, several TE families showed enrichment within EHMT2 peaks (Fig. S2J), including LTR8B, which also showed transcriptional upregulation in PEO1-R cells treated with combination therapy (Fig. 2A, 2K, Fig. S2K). These results show that inhibition of EHMT results in widespread epigenetic remodeling, including reactivation of EHMT-bound TEs (Fig. 2J-K).

Inhibition of epigenetic modifiers that represses transcription such as SETDB1 and DNA methylation has been linked to transcriptional reactivation of TEs, which can form immunostimulatory dsRNA that can subsequently induce an interferon response through a viral mimicry response^25, 26^. To determine if there is increased dsRNA formation in response to EHMT1/2 inhibition, we performed immunofluorescence of dsRNA on PEO1-R cells treated with control (DMSO), a PARP inhibitor (2*μ*M Olaparib), an EHMT inhibitor (1*μ*M UNC0642), or combination (2*μ*M Olaparib and 1*μ*M UNC0642) (Fig. 2L). No primary antibody and Decitabine (DAC) were used as negative and positive controls, respectively (Fig. 2K, 2M). Quantification showed that single UNC0642 and combination treatment significantly increased formation of dsRNA (Fig. 2M). Overall, these results showed that combinatory EHMT and PARP inhibition causes increased dsRNA formation, which may be due to transcriptional reactivation of multiple TE families that are repressed by EHMT1 and EHMT2.

### Response to combination therapy is partially dependent on RIGI and MDA5 *in vitro*

Since dsRNA was observed to be upregulated by combinatory EHMT and PARP inhibition, we asked whether intracellular sensors of dsRNA were responsible for triggering an interferon signaling cascade in combinatory EHMT and PARP inhibition therapy. Both RIGI and MDA5 were upregulated at the transcript level in PEO1-R cells treated with combination therapy (Fig. 3A-B). Single Olaparib and UNC0642 also led to increased transcription of RIGI and MDA5, respectively. To determine the contribution of RIGI and MDA5 to PARPi-resistant cell biology, we used CRISPRi to stably silence RIGI and MDA5 in PEO1-R cells and achieved a 45-65% reduction in transcript levels in both silenced cell lines (RIGI-knockdown/RIGI-KD, MDA5-knockdown/MDA5-KD) compared to our GFP control cells (Fig. 3C-D). Reduction of either RIGI or MDA5 did not affect overall cell growth compared to control cells (Fig. 3E). However, upon either single Olaparib or combination therapy, RIGI-KD cells showed increased viability compared to control cells, while MDA5-KD cells attenuated the cell growth defect observed in GFP control cells when treated with either single UNC0642 or combination therapy (Fig. 3F).

**Figure 3:**
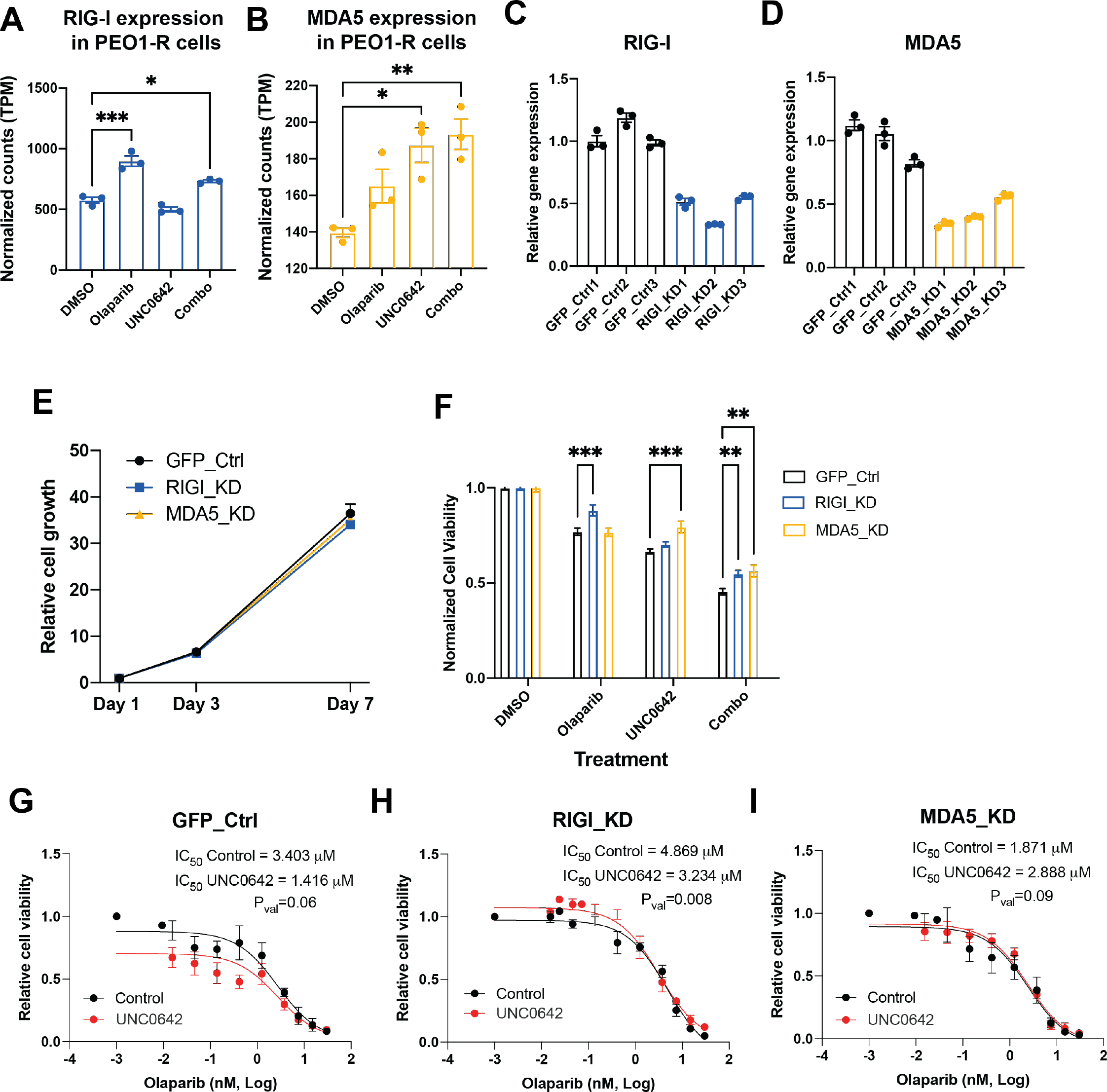
Response to combination therapy is partially dependent on RIGI and MDA5 *in vitro*. **A-B)** Gene expression of RIGI (A) and MDA5 (B) after PEO1-R cells were treated with DMSO, Olaparib, UNC0642, or Olaparib/UNC0642 for 72 hours. **C-D)** Gene expression of RIGI (C) and MDA5 (D) in PEO1-R cells expressing dCAS9 and gRNA targeting GFP (C-D), RIGI (C), and MDA5 (D). **E)** Cell growth of GFP control, RIGI-KD, and MDA5-KD cells measured by cell titer glo assay. **F)** Cell viability of GFP control, RIGI-KD, and MDA5-KD cells after being treated with DMSO, Olaparib, UNC0642, or Olaparib/UNC0642 for 7 days. **G-I)** GFP control (G), RIGI-KD (H), and MDA5-KD (I) cells treated with control (DMSO, black lines) or an EHMT inhibitor (UNC0642, red lines) and increasing doses of Olaparib for 5 days. Error bars, SEM. Statistical test, multi-comparison ANOVA and non-linear regression. *p<0.05, **p<0.01, ***p<0.001.

We have shown in Watson et al^18^ that UNC0642 can increase sensitivity to Olaparib in PARPi resistant PEO1-R cells. To determine if PARPi resensitization was dependent on RIGI or MDA5, Olaparib dose response assays were performed on PEO1-R cells with and without UNC0642 treatment. Our results show that co-treatment with UNC0642 trended towards resensitizing GFP control cells to Olaparib (p-value=0.06) with decrease in IC50 from 3.4μM in cells without UNC0642 co-treatment to 1.4μM in cells with UNC0642 co-treatment (Fig. 3G). While RIGI-KD cells did show a resensitization effect (Fig. 3H), MDA5-KD cells attenuated the resensitization effect observed in GFP control and RIGI-KD cells with UNC0642 co-treatment (Fig. 3I). Altogether, our results indicate that silencing RIGI or MDA5 does not affect PARPi-resistant cells at baseline but does inhibit proliferation in response to combinatory PARP and EHMT inhibition. These findings indicate that the anti-proliferative effects of combinatory PARP and EHMT inhibition is partially dependent on RIGI and MDA5, implicating dsRNA-mediated interferon signaling as key mechanism underlying the anti-tumor effect of combined PARP and EHMT inhibition.

### Immune intact PARP inhibitor resistant model

Given that combinatory EHMT and PARP inhibition drives activation of several immune signaling pathways including the interferon responses, we next investigated how combination therapy affects anti-tumor immunity in an intact immune *in vivo* model of PARPi resistant ovarian cancer.

First, we defined the expression of EHMT1 and EHMT2 across tumor microenvironment cell types by analyzing a single-cell RNA-seq dataset from five tumor patient samples^36^. We examined tumor cells (PAX8+), macrophages (CD68+), fibroblasts (FAP+), and T cells (CD3+) and found that EHMT expression was largely restricted to tumor cells with a portion of EHMT-expressing fibroblasts and macrophages (Fig. S3). Thus, to interrogate the activation of immune pathways *in vivo*, we exposed ID8 Trp53-/-, Brca2-/- syngeneic mouse cells developed by Walton et al^37^ to increasing doses of Olaparib to establish a PARPi-resistant (ID8-R) strain of a mouse ovarian cancer cell model. ID8-R cells have a 17-fold increase in Olaparib IC50 compared to their sensitive ID8 counterparts (Fig. 4A).

**Figure 4:**
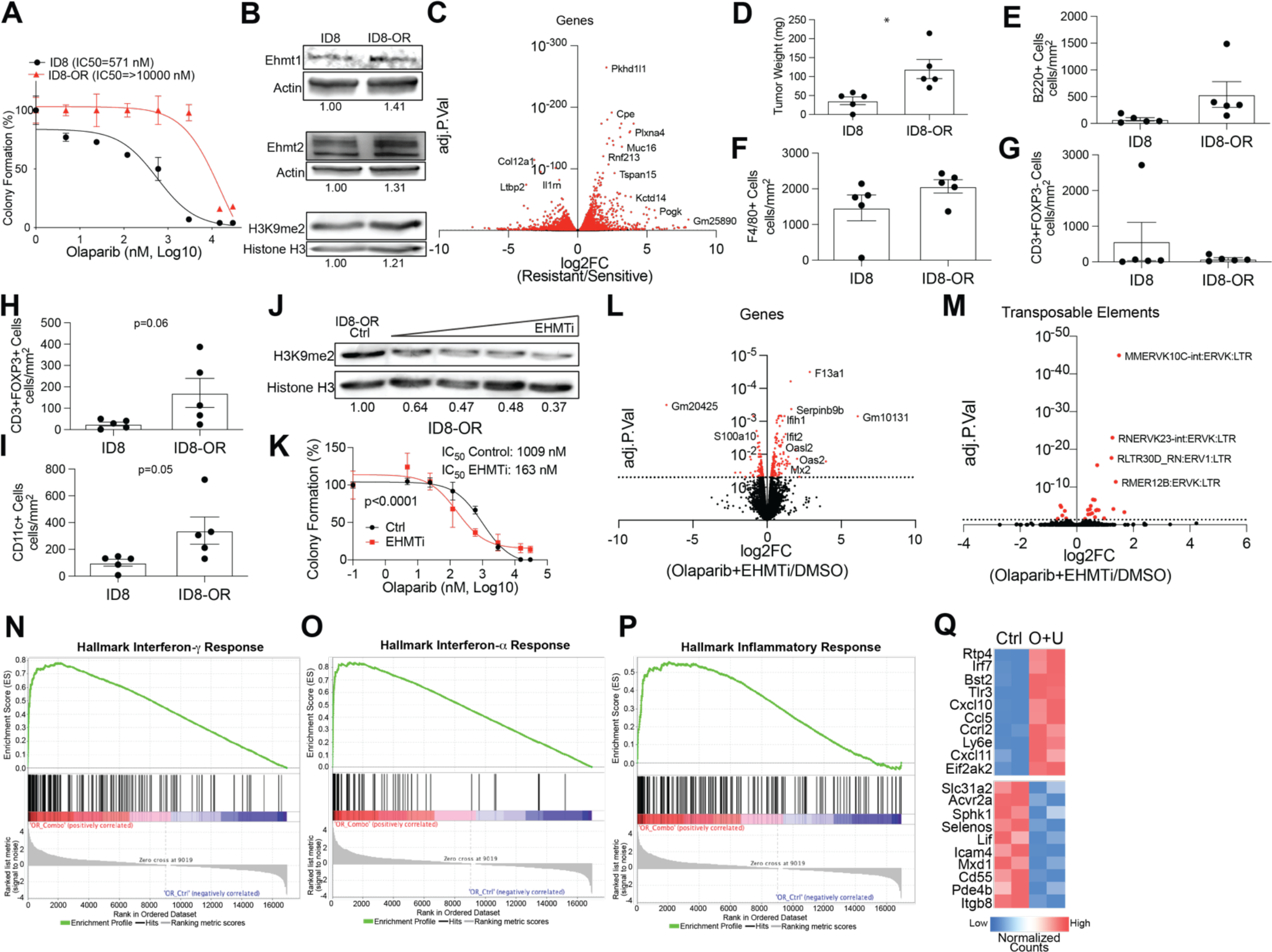
An immunogenic model of PARP inhibitor resistant ovarian cancer. **A)** ID8 Trp53-/-, Brca2-/- sensitive (ID8) and Olaparib resistant (ID8-R, red line) treated with increasing doses of Olaparib. **B)** Protein from ID8 and ID8-R used for immunoblot against EHMT1, EHMT2, and H3K9me2. Loading controls, Actin and Histone H3. **C)** Volcano plot of ID8-R transcriptome compared to ID8 cells. **D)** *In vivo* evaluation of tumor burden 35 days after cell implantation. ID8 and ID8-R tumors were fixed and used for evaluation via multispectral IHC. The density of different cell types is graphed. **E)** B cells via B220+ cells, **F)** macrophages via F4/80+ cells, **G)** non-T regulatory cells via CD3+, FOXP3-, **H)** T regulatory cells via CD3+, FOXP3+, **I)** dendritic cells via CD11c+. **J)** ID8-R cells treated with increasing doses of UNC0642 and protein blotted against H3K9me2. Loading control, Histone H3. **K)** ID8-R cells treated with control (DMSO, black lines) or EHMT inhibitor (EZM8266, red lines) and increasing doses of Olaparib for 12 days. **L)** Volcano plot of differentially regulated genes in ID8-R cells treated with Olaparib/UNC0642 compared to vehicle control (red dots, adj. p-value<0.05). **M)** Volcano plot of differentially regulated transposable elements in ID8-R cells treated with Olaparib/UNC0642 compared to vehicle control (red dots, adj. p-value<0.05). Hallmark geneset enrichment of ID8-R cells treated with combination for **N)** Interferon-*γ* response, **O)** Interferon-*α*response, and **P)** Inflammatory response. **Q)** Heatmap of significantly differentially regulated genes in control (Ctrl) and Olaparib/UNC0642 (O+U) in the inflammatory pathway. Error bars, SEM. Statistical test, unpaired t-test with Welch’s correction and Log-rank. *p<0.05.

Similar to our human models of PARPi-resistant ovarian cancer cells, ID8-R cells have elevated EHMT1 and 2 gene and protein expression that corresponded to an increase in H3K9me2 (Fig. 4B). Compared to the ID8 cells, the ID8-R cells have a significantly different transcriptome that includes elevated expression of the ovarian cancer tumor marker, Muc16 (Fig. 4C). To understand the difference in the tumors generated from the ID8-R cells compared to the ID8 cells, cells were injected into immune intact female C57Bl6 mice and allowed to grow untreated. Consistent with clinical progression of recurrent and therapy-resistant disease, the therapy resistant ID8-R tumors were more aggressive as indicated by tumor burden (Fig. 4D). Next, multispectral immunohistochemistry (mIHC) was used to examine the tumor composition from the ID8 and ID8-R cells. The density of tumor cells (WT1+), macrophages (F4/80+), B-cells (B220+), non-regulatory T-cells (CD3+, FOXP3-), regulatory T cells (CD3+, FOXP3+), dendritic cells (CD11c+), and neutrophils (Ly6G+) were calculated from the ID8 and ID8-R tumors (Fig. 4E-I). Neutrophils (Ly6G+) were not detected within any tumor evaluated. Notably, both regulatory T cells and macrophages show a trend toward enrichment in the ID8-R cells (Fig.4F,H). Dendritic cells (CD11c+) cells were the only significantly enriched cell type in the ID8-R tumors compared to ID8 tumors (Fig. 4I, p=0.05). These data demonstrate the ID8-R tumors are aggressive with differential tumor immune compositions that suggest an immune suppressed tumor microenvironment.

Next, we examined how ID8-R cells respond to the combination of EHMT and PARP inhibition *in vitro*. Following treatment of ID8-R cells with different concentration of an EHMT inhibitor, H3K9me2 is depleted and the sensitivity to PARP inhibition is increased in a dose-dependent fashion (Fig. 4J-K). Similar to the human-derived PARPi resistant cells, the combination of EHMT and PARP inhibition *in vitro* promotes differential gene and TE family-level expression (Fig. 4L-M). Consistently, the combination treatment led to the activation of several immune related pathways, including interferon-*γ* response, interferon-*α* response, and inflammatory response (Fig. 4N-P). Examining the inflammatory response, several immune factors and cytokines were noted to be elevated following the combinatory treatment (Fig. 4Q). These factors are essential for T cell recruitment into the microenvironment and higher expression of CCL5, CXCL10, and CXCL11 in human ovarian cancer tumors are noted to convey a 25-month improved overall survival (Fig. 1M, Low Expression=41.6 months versus High Expression=66.4, Logrank p<0.0001). These data show that the murine ID8-R model similarly responds to EHMT inhibition through gene regulation, TE transcriptional activation, and immune pathway activation as human PARPi-resistant cell models.

### Targeting EHMT in an immune intact PARPi resistant *in vivo* model

Given the activation of immune related pathways, we next evaluated the anti-tumor response and immune microenvironment following combinatory EHMT and PARP inhibition *in vivo* (Fig. 5A). GFP-luciferase-tagged ID8-R tumor bearing mice were randomized and treated with vehicle control, Olaparib, EHMT inhibitor (EZM8266), or combination (Fig. 5A and S4A) (experiment 1). EZM8266 is a novel, orally available EHMT1/2 inhibitor that has superior oral bioavailability compared to UNC0642 *in vivo*. Tumor progression was monitored via IVIS imaging over the course of treatment. The Olaparib-treated ID8-R tumors demonstrated resistance *in vivo*, growing at a similar rate as control tumors (Fig. 5B). Notably, mice treated with EZM8266 alone showed reduced total photon flux after 14 days, suggesting tumor regression and a potential anti-tumor immune response. Upon necropsy after 28 days of treatment, the omental weight, number of dissemination sites, and solid tumor nodule weight were all significantly reduced across the different treatment groups compared to vehicle treated tumors (Fig. 5C-E). While the tumor burden was reduced following Olaparib treatments, both single EZM8266 and combinatory EZM8266/Olaparib treated mice had significantly fewer dissemination sites and lower solid tumor weight compared to mice treated with Olaparib alone. At the time of necropsy, tumor-bearing mice underwent a Complete Blood Cell count panel to evaluate drug toxicity. We did not observe any overt signs of drug toxicity based on Complete Blood Cell count (Fig. S4B-D) and body weight (Fig. S4E). In an independent *in vivo* study using the ID8-R (experiment 2), the EZM8266 anti-tumor response was noted to be dose-dependent (Fig. S4F-J).

**Figure 5:**
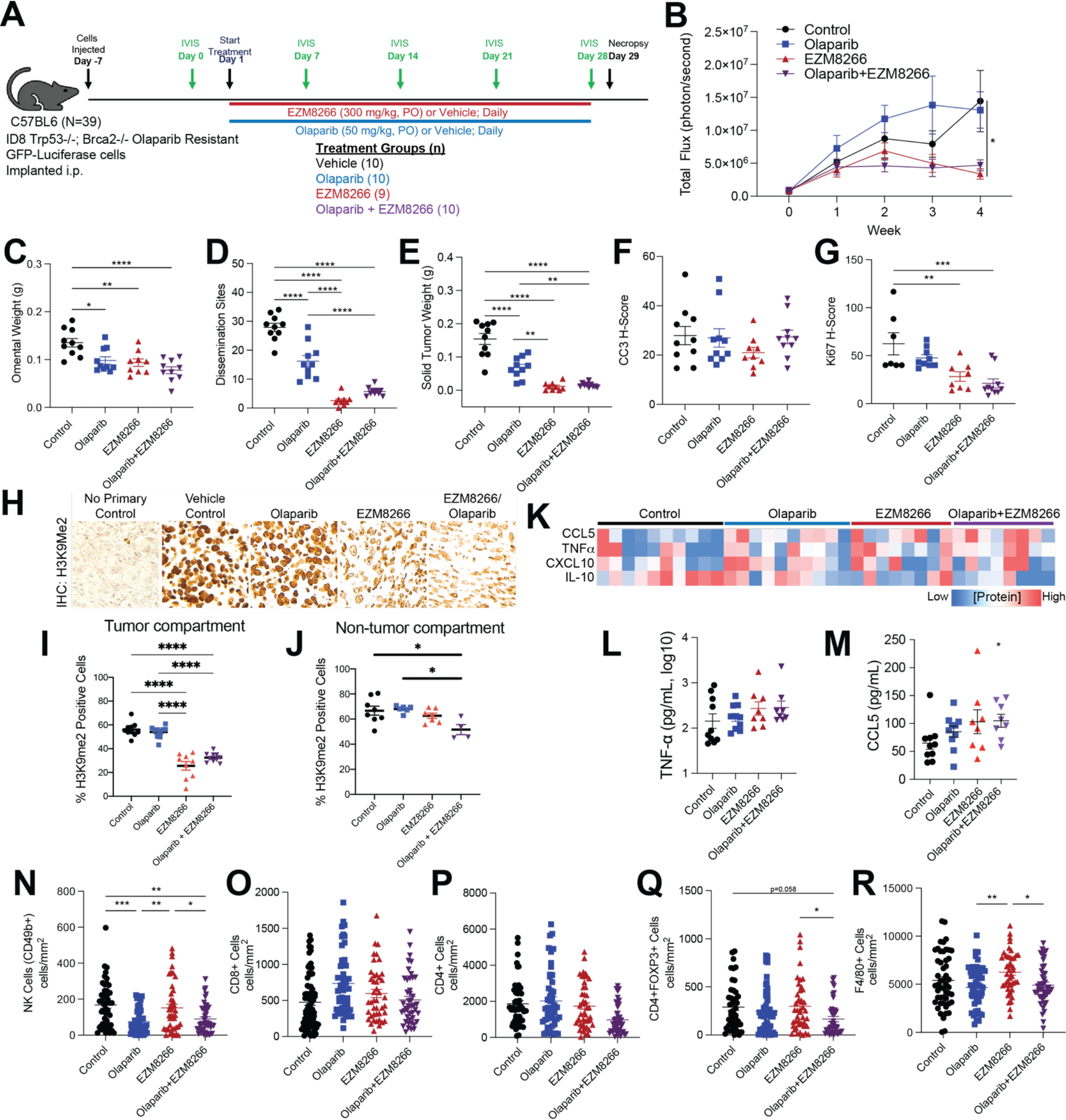
EHMT inhibition significantly reduces tumor progression in an Olaparib resistant ovarian cancer syngeneic mouse model. **A)** Study design and timeline. Population size is indicated by treatment group (n= 9 or 10). **B)** Tumor progression monitored via total flux (photons/sec). **C)** Omental weight. **D)** Number of tumor dissemination sites detected. **E)** Weight of solid tumors. **F)** Histological score (H-score) of formalin fixed tumor used for IHC against cleaved caspase 3 (CC3). Histological score (H-score) of formalin fixed tumor used for IHC against Ki67. **H)** IHC against H3K9me2. **I)** Percent H3K9me2 positive cells in tumor compartment. **J)** Percent H3K9me2 positive cells in non-tumor compartment. **K)** Heatmap of multiplex ELISA for indicated cytokines in ascites fluid collected from treated tumor-bearing mice. **L)** Graphical data of TNF-*α* concentration. **M)** Graphical data of CCL5. Tumors from treated mice were used for mIHC and 5 ROI were selected from each tumor. Tumor infiltration of **(N)** NK cells, **(O)** CD8+ cells, **(P)** CD4+ cells, **(Q)** CD4+FOXP3+ cells, and **(R)** F4/80+ cells. Error bars, SEM. Statistical test, mixed model effect (B) and multicomparison ANOVA with Tukey correction (C-G, J-P). *p<0.05, **p<0.01, ***p<0.001, ****p<0.0001.

These studies show EHMT inhibition is a safe and sufficient approach to induce regression of PARPi-resistant tumors.

Next, downstream IHC was performed to characterize apoptosis (cleaved caspase 3) and proliferation (Ki67) in tumor-bearing mice. Neither single nor combination-treated tumors showed a significant difference from vehicle control in apoptosis (Fig. 5F). In contrast, EZM8266 and EZM8266/Olaparib tumors had significantly lower Ki67 H-scores than vehicle control, indicative of a treatment induced anti-proliferation response (Fig. 5G). EHMT1/2 catalyze the H3K9me2 modification, thus EZM8266 and combination-treated tumors were confirmed to have a reduction in the H3K9me2 modification (Fig. 5H). While a mild reduction of H3K9me2 was observed in non-tumor compartment with combination therapy, the tumor compartment had drastically more reduction of H3K9me2 with single and combinatory EHMT inhibition (Fig. 5I-J), suggesting specificity of the EHMT targeting.

In addition to solid tumors, ascites fluid and cells can contribute markedly to patient disease burden. Based on RNA-seq data, combinatory EHMT and PARP inhibition induce an inflammatory response. We accordingly interrogated the protein concentration of pro-inflammatory (CCL5, CXCL10, TNFα) and anti-inflammatory (IL-10) cytokines in the ascites fluid of tumor bearing animals using multiplex ELISA. Both single EZM8266 and combinatory EZM8266/Olaparib led to an increase in pro-inflammatory cytokines compared to control treated mice (Fig. 5K-M). Conversely, IL-10 was downregulated in the EZM8266 and EZM8266/Olaparib treated mice (Fig. 5K). CCL5 was significantly upregulated in the EZM8266/Olaparib treated tumors compared to control (Fig. 5M) within the collected ascites fluid.

Multispectral IHC on tumor sections was completed to interrogate differential tumor immune cell composition. Compared to control, CD49b^+^ (NK cells) were depleted in tumors treated with either Olaparib alone or EZM8266/Olaparib (Fig. 5N) and unchanged in EZM8266-only treatment. Also, compared to control treatment, there was not a significant difference in tumor-associated CD8^+^ and CD4^+^ T cells (Fig. 5O-P). However, in the EZM8266/Olaparib combination there was a significant reduction in CD4^+^FOXP3^+^ T regulatory cells compared to control or EZM8266 alone. F4/80^+^ macrophages were enriched in EZM8266 tumors compared to control and EZM8266/Olaparib combination (Fig. 5Q-R).

In summary, the use of EZM8266 alone or in combination with Olaparib in the ID8-R mouse model effectively reduces disease progression and primary tumor burden, and sharply curtails metastasis. EZM8266 also effectively reduces H3K9me2 and Ki67 in tumors and alters the cytokine milieu of ascites fluid. Further, EZM8266 and combination-treated tumors exhibit a depletion of regulatory T cells, emphasizing that EHMT inhibition is sufficient to drive tumor regression and remodeling of the tumor immune microenvironment.

### EHMT inhibition induces a T cell dependent anti-tumor response

Across both human and mouse models of PARP inhibitor resistance, EHMT and PARP inhibition led to increased activation of several immune response pathways. Several pro-inflammatory cytokines were noted to be increased following EHMT and PARP inhibition, including CCL5, CXCL10, and CXCL11. These cytokines have established roles in recruiting T cells into the tumor microenvironment^38^. Deconvoluting RNA-seq data from ovarian cancer TCGA tumors revealed significant positive correlations between *CCL5*, *CXCL10*, and *CXCL11* mRNA expression and CD8+ T cell tumor infiltration (Fig. 6A-C). To determine the acute T cell response of PARP and EHMT inhibition, *ex vivo* cultures of primary tumors were treated with control (DMSO), a PARP inhibitor (1*μ*M Olaparib), an EHMT inhibitor (1*μ*M UNC0642), or combination (1*μ*M Olaparib and 1*μ*M UNC0642). Immune cell composition and activation were evaluated in tumor sections using mIHC. In three independent tumors, single PARP and EHMT inhibition did not significantly alter the regulatory T cell (CD4+/FOXP3+) proportion. However, combinatory PARP and EHMT inhibition significantly elevated granzyme B+ T cells compared to the vehicle control (Fig. 6D-E), suggesting activation of effector T cell function.

**Figure 6:**
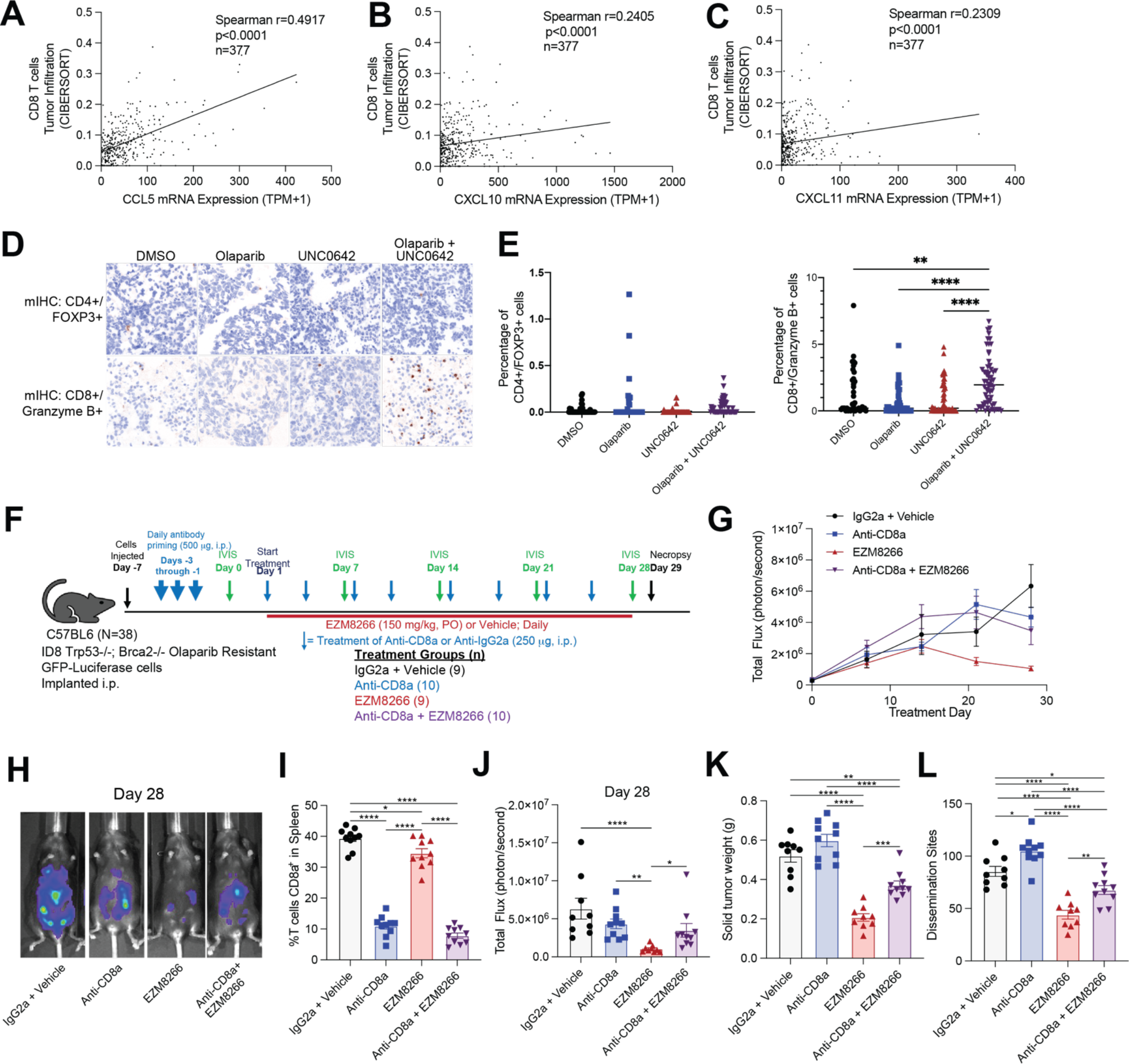
Anti-tumor response of EHMT inhibition is dependent on CD8 T cells. CIBERSORT analysis of ovarian cancer tumors (TCGA PanCancer Atlas) examining CD8+ T cell infiltration compared to **A)** *CCL5*, **B)** *CXCL10*, and **C)** *CXCL11* mRNA expression. **D)** Primary ovarian cancer tumors treated with EHMT inhibitor (UNC0462) and/or Olaparib. Tumor sections were fixed and analyzed via multispectral IHC. Representative images of tumors for regulatory T cells (CD4+/FOXP3+) and effector T-cells (CD8+/Granzyme B+). **E)** Quantification of regulatory T cells (CD4+/FOXP3+) and effector T-cells (CD8+/Granzyme B+). **F)** Study design and population size indicated next to treatment (n=9 or 10/group). G) Tumor progression measured via total flux. **H)** Representative images of luminescence in tumor-bearing mice at the end of study for each treatment group. **I)** Percentage of splenic CD8a+ T cells at the end of study. **J)** Tumor burden measured as total flux on Day 28. **K)** Solid tumor weight. **L)** Number of disseminated tumors. Error bars, SEM. Statistical test, multi-comparison ANOVA with Tukey correction. *p<0.05, **p<0.01, ***p<0.001, ****p<0.0001.

To further test the dependence on T cells of the anti-tumor response observed *in vivo*, in experiment 3, CD8 T cells were either depleted via CD8 neutralizing antibody or left intact (isotype control treatment) in ID8-R tumor bearing mice. CD8 T cell depleted mice were then treated with biweekly CD8 neutralizing antibody and/or EZM8266, and CD8 T cell-intact mice were treated with biweekly isotype control antibody and/or EZM8266 (Fig. 6F and S5A-C). Strikingly, after 28 days of treatment the depletion of CD8 T cells attenuated the EZM8266-induced anti-tumor response (Fig. 6G-H). Also, after 28 days of treatment, CD8 T cells remained depleted based on splenic CD8+ T cells analysis (Fig. 6I). In EZM8266 treated mice, CD8 T cell depletion partially attenuated the total flux, tumor weight, and number of dissemination sites compared to isotype/EZM8266 treated mice (Fig. 6J-L). These data demonstrate that EHMT inhibition promotes the activation of T cells, and that the anti-tumor response is partially dependent on the presence of CD8+ T cells.

### Combination therapy may contribute to T cell exhaustion and dsRNA-induced interferon response *ex vivo*

To further interrogate the mechanism of action in primary tumors, we used mIHC to interrogate several mechanisms including apoptosis (cleaved caspase 3), proliferation (Ki67), DNA damage (*γ*H2AX), T cell exhaustion (CD3, LAG3, PD1, PDL1), and downstream components of the RIGI/MDA5 pathway (MAVS, IRF9, pSTAT1) in patient-derived, therapy-naïve, *ex vivo* tissues treated with control (DMSO), a PARP inhibitor (2*μ*M Olaparib), an EHMT inhibitor (1*μ*M UNC0642), or combination (2*μ*M Olaparib and 1*μ*M UNC0642). Similar to our ID8-R model, reduction of H3K9me2 was mainly confined to the tumor compartment (Fig. 7A-B). Apoptosis was significantly increased in tissues treated with combination therapy compared to Olaparib treatment (Fig. 7C), and DNA damage was significantly increased with single UNC0642 treatment compared to single Olaparib treatment (Fig. 7D). Proliferation was not significantly changed with any of the treatments (Fig. 7E). PDL1 expression on tumor cells was significantly increased with combination treatment, while LAG3 and PD1 expression on CD3 cells were repressed with Olaparib treatment compared to control and single UNC0642 treatment (Fig. 7D-F). Lastly, downstream components of the RIGI/MDA5 pathway were mostly unperturbed by treatments overall and in H3K9me2+ cells (Fig. 7J-L, Fig. S6A-C). However, in H3K9me2- cells, MAVS was increased by single UNC0642 and combination treatment compared to single Olaparib treatment (Fig. 7M); pSTAT1 was increased by single UNC0642 and combination therapy compared to DMSO (Fig. 7N); and IRF9 was increased by single UNC0642 treatment compared to single Olaparib treatment (Fig. 7O). Altogether, these mIHC studies indicate that single EHMT inhibition, or combinatory PARP and EHMT inhibition contribute to T cell exhaustion and dsRNA-induced interferon response.

**Figure 7:**
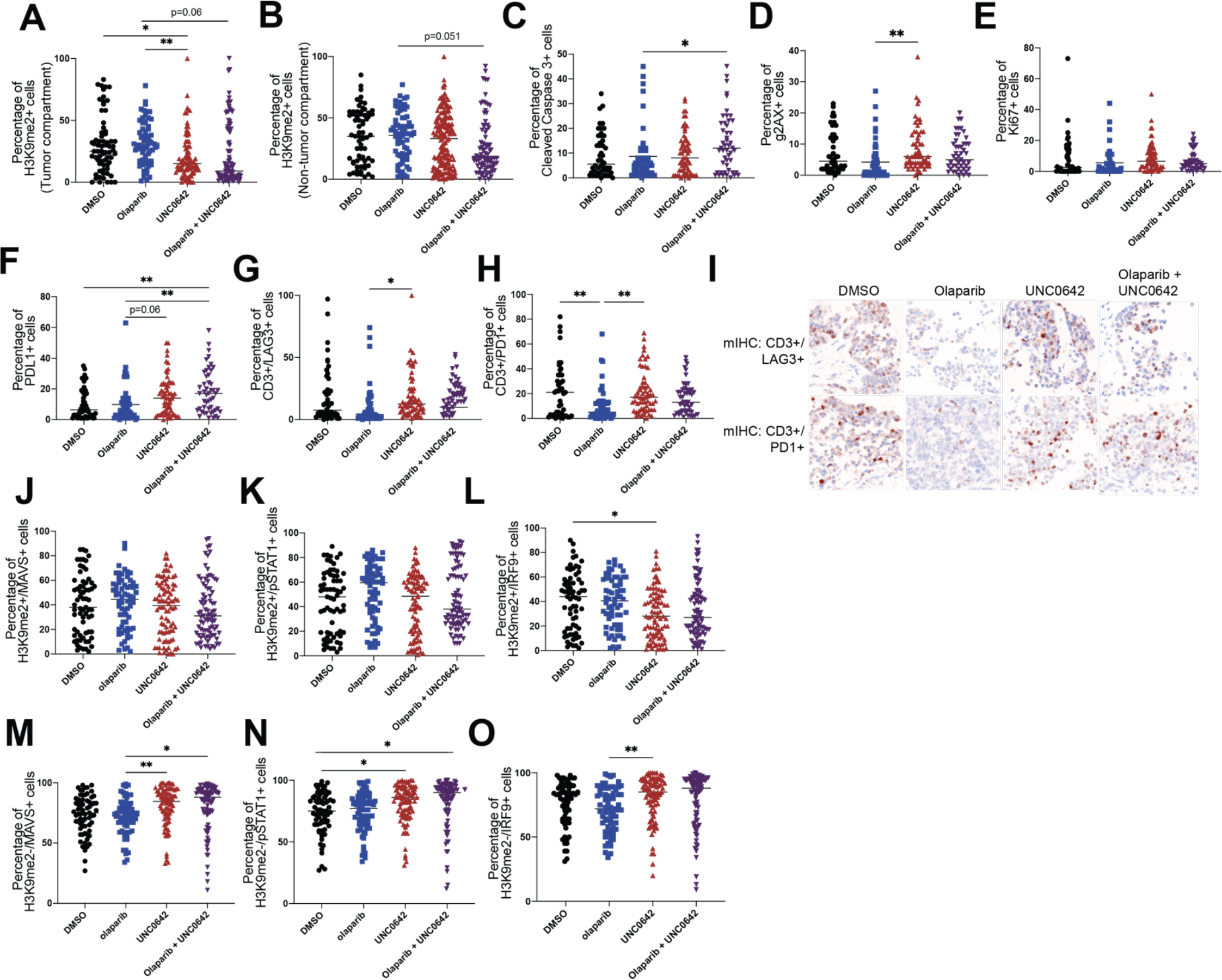
Combination therapy may contribute to T cell exhaustion and dsRNA-induced interferon response *ex vivo*. Primary ovarian cancer tumors treated with EHMT inhibitor (UNC0462) and/or Olaparib. Tumor sections were fixed and analyzed via multispectral IHC. Quantification of **A)** H3K9me2 positive cells in tumor compartment, **B)** H3K9me2 positive cells in non-tumor compartment, **C)** apoptosis (cleaved caspase 3+), **D)** DNA damage (gH2AX+), **E)** proliferation (Ki67+), **F)** PDL1, **G-**T cell exhaustion (CD3+/LAG3+ and CD3+/PD1+). **I)** Representative images of tumors for T cell exhaustion (CD3+/LAG3+ and CD3+/PD1+). **J-L)** Quantification of downstream RIGI/MDA5 proteins (MAVS+, pSTAT1+, IRF9+) in H3K9me2+ cells and in **M-O)** H3K9me2- cells. Error bars, SEM. Statistical test, multi-comparison ANOVA with Tukey correction. *p<0.05, **p<0.01, ***p<0.001, ****p<0.0001.

## DISCUSSION

Since its discovery in 2005, several PARP inhibitors (e.g. olaparib, rucaparib, and niraparib) have been FDA approved for the treatment of ovarian cancer for both homologous recombination deficient and proficient genotypes^3–5^. PARP inhibitors are also FDA approved for breast, prostate, and pancreatic cancer, and are currently under trial for several other cancer types^8–10^. However, like many other therapies, cancer cells eventually develop resistance to PARP inhibitors^12^. Due to its widespread and continually growing use, development of alternative therapies for use in PARP inhibitor resistant tumors is vital for improving patient outcomes.

Our lab has shown that PARP inhibitor resistant ovarian cancer cells have increased levels of EHMT1/2 and the repressive histone mark that these two enzymes deposit, H3K9me2^18^. Additionally, reducing EHMT1/2 expression in PARPi-resistant cell lines resensitized the cells back to PARPi treatment^18^. In this study, we used a combination of cellular biology, bioinformatics, and multiple *in vitro* and *in vivo* models to determine the mechanism and efficacy of combinatory EHMT and PARP inhibition in the treatment of PARPi-resistant ovarian cancer. We discovered that, on an organism level, single EHMT and combinatory PARP and EHMT inhibition was effective in reducing PARPi-resistant ovarian tumors and that this reduction in tumor growth is dependent on CD8 T cells. Our cellular and mechanistic studies suggest that this dependency and involvement of CD8 T cells may be a result of increased interferon response induced by EHMT-mediated TE reactivation. In line with other studies of epigenetic therapies, we found that inhibiting EHMT results in increased epigenetic and transcriptional activation of TEs, as well as increased dsRNA. We confirmed that these dsRNA are detected by RIGI and MDA5, triggering an interferon response that causes secretion of cytokines (e.g. CCL5) that activate multiple immune signaling pathways and recruit T cells.

Our RNA-seq analysis of several PARPi-resistant cells and therapy-naïve, patient-derived *ex vivo* tissues revealed that that combinatory EHMT and PARP inhibition robustly induces several immune signaling pathways, mainly interferon-*α* and interferon-*γ* signaling pathways. In addition, transposable element (TE) transcripts were also consistently upregulated in cells treated with combinatory EHMT and PARP inhibition. This is in line with current literature on epigenetic therapy and TE reactivation. Roulois et al^26^ and Chiappinelli et al^25^ have shown that inhibition of DNA methyltransferase (DMNT) and reduction of its repressive mark, DNA methylation, can uncover and reactivate TEs. Reactivation of TEs can increase immunostimulatory dsRNA formation, trigger an interferon response, and promote an anti-tumor response in ovarian and colorectal cancer cells^25, 26^. While we were able to confirm the involvement of dsRNA innate immune sensors and we detected dsRNA upregulation by immunofluorescence, our study did not directly test whether TEs were the primary source of immunostimulatory RNAs, though the immunostimulatory effect of many TE-derived RNAs including from LINE1 elements and Alus has been demonstrated by recent studies^25, 26, 39^.

It is interesting that in our studies, both PARP and EHMT inhibition is required to induce TE levels and interferon signaling. EHMT inhibition can reactivate TEs in ovarian cancer cells but has only been shown in combination with DMNT inhibition^14^, which broadly derepresses the genome regardless of cell type. To our knowledge, PARP1/2 has not been shown to regulate TEs. Our observations can be explained by the ability of PARP inhibitor’s ability to induce dsDNA formation and stimulate interferon signaling via the cGAS-STING pathway, compounding with additional stimulation of interferon signaling from EHMT inhibition.

Our functional experiments and epigenomic profiling studies support a role for EHMT1/2 in TE regulation. In contrast to the TE reactivation observed with combinatory EHMT and PARP inhibition, we observed a repression of TEs in cells when EHMT1 or EHMT2 was overexpressed. Next, we show that EHMT inhibition not only affects EHMT1/2 catalytic activity (dimethylation of H3K9), but also chromatin targeting of EHMT2. Additionally, we show that at the same sites in which EHMT2 localizes, there is an increase in open chromatin and enhancer marks when treated with combinatory EHMT and PARP inhibitors, suggesting that EHMT2 maintains repressive chromatin structure. Lastly, EHMT2 localizes to both protein-coding genes and TE loci, providing evidence that EHMT2 directly interacts with TEs. This, in combination with the repression of TEs in EHMT1 and EHMT2 overexpressed cells, suggest that EHMT1/2 play an important role in the regulation of TE transcript expression. Derepression of TEs by EHMT1/2 inhibition could also have other effects beyond transcription, including altering nearby gene expression by regulating TE-derived enhancers and promoters^40, 41^.

We postulate that the interferon response to combination therapy was driven by dsRNA, which is supported through knock-down studies of intracellular sensors of dsRNA, RIGI and MDA5. Based on our results, RIGI regulates Olaparib-driven effects while MDA5 regulates UNC0642-driven effects. This would align with the recently appreciated distinct RNA binding features and signaling pathways of RIGI and MDA5^42^. RIGI preferentially binds to shorter dsRNA with tri-phosphorylation at its 5’ end^42^, while MDA5 binds to longer dsRNAs. RIGI has also been shown to bind to RNA:DNA hybrids^43^, while MDA5 is shown to be more interactive with TEs^39, 44^. Depletion of PARP1 causes genome instability which can produce R-loops (DNA-RNA hybrids)^45^, while UNC0642 has been shown here and in other studies to regulate TE transcript expression^27, 45, 46^. Taken together, we hypothesize that RIGI and MDA5 are both contributing to the interferon response by binding to DNA:RNA hybrids formed by PARPi-induced genome instability and TE-derived dsRNA reactivated by UNC0642, respectively. One limitation from this study, however, is that even though we show correlation between TE and dsRNA, we have not directly shown that TEs are the primary source of immunostimulatory dsRNA. Additionally, we did not examine the contributions of other dsRNA sensors, such as LGP2^47^.

Given the partial dependency on RIGI and MDA5 and the robust interferon and immune signaling responses, we decided to utilize an *in vivo* model that contains an intact immune system. To that end, we used the ID8 TP53-/-, BRCA2-/- syngeneic cells developed by Walton et al to create an Olaparib-resistant cell syngeneic model^37^. Characterization of these cells and *in vivo* model showed that we were successful in achieving PARPi-resistance. To our knowledge, this is the first PARPi-resistant syngeneic *in vivo* model. The immune system has emerged as an important component in the treatment of cancer, and countless combinatory immunotherapies are emerging, therefore a relevant immune-intact *in vivo* model is necessary to study potential therapies^48^. Some caveats of our *in vivo* model include the GFP-luciferase tag on the tumor cells which are immunogenic. However, our control cells also have a luciferase tag and should correct for baseline activation of immune cells. Another caveat is that the therapy in mouse cells may differ from that in human cells. However, our characterization showed that these mouse ID8-R cells and human PARPi-resistant cells respond similarly to single and combinatory PARP and EHMT inhibition. Since these cell autologous responses mirror each other, we believe that this ID8-R, immune-intact, syngeneic model is appropriate to study the mechanisms of combinatory PARP and EHMT inhibition in the treatment of PARPi-resistant ovarian cancer. We also believe this model could be used to test other therapies after verifying that the responses in human cell lines are recapitulated in this ID8-R cell line.

When we tested single and combinatory PARP and EHMT inhibition in our ID8-R syngeneic model, we found that all treatments significantly reduced tumor burden. However, single EHMT and combinatory EHMT and PARP inhibition had the greatest reduction in tumor burden. This effect may be due to several mechanisms. Ki67 staining was decreased, indicating a decrease in tumor cell proliferation. Additionally, T regulatory cells were decreased and several cytokines (CCL5, CXCL10, TNFα) were increased, suggesting a remodeling of the tumor immune microenvironment. Indeed, further characterization of the tumor microenvironment using multispectral IHC and functional T cell markers, we observed that cytotoxic CD8 T cell activity (via Granzyme B) was increased in *ex vivo* tissues treated with combinatory EHMT and PARP inhibition. Most importantly, we showed that the tumor reduction driven by EHMT inhibition is dependent on CD8 T cells. These results have huge implications for combining EHMT inhibition with immunotherapies, especially when our mIHC analyses showed differential states of T cell exhaustion with different treatments in our *ex vivo* tissues. Epigenetic therapies have been combined with immunotherapy and show efficacies in several cancer models^21^ suggesting that combinatory EHMT inhibition and immunotherapy may have similar benefits. Lastly, even though we have strong correlative data between TE and dsRNA *in vitro* and downstream components of the RIGI/MDA5 pathway and combination therapy *ex vivo*, we have not done functional studies to test the necessity and sufficiency of dsRNA-induced interferon response *in vivo*.

Unlike our cell autologous model, however, single EHMT inhibition alone can promote this putative anti-tumor immune response. Whether or not PARP inhibition is required for further improved response is still to be determined. A survival study can perhaps show a long-term benefit of combinatory treatment versus single. Another interesting observation from our animal models is that ablation of CD8 T cells only partially rescued the tumor reduction phenotype, suggesting other EHMT- driven mechanisms. This is consistent with current literature on EHMT in cancer and therapy resistance. High EHMT1/2 expression is associated with poorer survival outcomes and therapy resistance, though the mechanisms seem to be context-dependent^20^. In the context of PARPi-resistant ovarian cancer, our data suggest a large dependency on CD8 T cells and perhaps EHMT- mediated changes in gene regulation that we have yet to functionally validate.

Overall, we show that combinatory EHMT and PARP inhibition is effective in treating PARPi-resistant ovarian cancer at the cellular and organism level. We presented evidence that the mechanism is CD8 T cell dependent and is strongly associated with reactivation of transposable elements. Though this study started because we found that PARPi-resistant cells have increased EHMT1/2 compared to non-resistant cells, ovarian tumor cells intrinsically have more EHMT1/2 expression that normal ovarian cells suggesting a possible efficacy in therapy-naive ovarian cancer. Additionally, based on specific upregulation of EHMT1/2 expression in other cancer types^20^, this combinatory or even single EHMT therapy may be effective in treating those cancers as well. Lastly, our finding that EHMT therapy stimulates the activation of CD8 T cells has major implications for combining EHMT inhibitors with immunotherapy such as immune checkpoint blockade.

## METHODS

### Cell Culture

PEO1 (RRID:CVCL_2686), Kuramochi (RRID:CVCL_1345), and OVCA420 (RRID:CVCL_3935) cell lines were provided through the Gynecologic Tumor and Fluid Bank (GTFB). ID8 cell line was kindly gifted from Dr. Iain McNeish lab (Imperial College London). All cells were authenticated via STR at The University of Arizona Genetics Core (RRID:SCR_012429). All cells undergo monthly mycoplasma testing using Sigma LookOut Mycoplasma Detection kit (Cat. #MP0035) and are only culture up to 20 passages or two months. Cells are cultured in RPMI1640 (Gibco) in 10% FBS (Phoenix Scientific Cat. #PS-100) and Penicillin Streptomycin (Cat. #SV30010) at 37C in 5% CO2. ID8 cells are cultured in DMEM (Gibco, Cat. #11995-065) supplemented with 4% FBS, Penicillin Streptomycin and 1x ITS (Gibco Cat. #41400-045).

### Ex vivo Culture

Primary ovarian cancer tumors (Table S1) were collected at The University of Colorado Cancer Center under the Colorado Multiple Institutional Review Board (COMIRB) Gynecologic Tumor and Fluid Bank protocol #07-935. All patients are consented under the GTFB protocol and collected specimens are deidentified. Tumors are collected by the Biorepository Shared Resource and delivered to the lab within one hour of surgical dissection. Tumors are sliced using a Krumdieck Tissue Slicer into 300-micron thick discs and placed in warmed culture media. Tumor sections treated for 72 hours, are either stored at -80C in RLT plus buffer (Qiagen, Cat. #74136) or fixed in 10% buffered formalin. RNA was extracted from tissues stored in the RLT buffer and used for RNA-sequencing. Fixed tissues were embedded in paraffin at the University of Colorado Histopathology Shared Resources.

### Cell culture and generation of EHMT1 and 2 overexpression

PEO1 cells were maintained in RPMI+ 10% heat inactivated FBS (Avantor, Cat. #10803-034) + 100 μg/mL Penicillin/Streptomycin (Lonza, Cat. #09-757F). EHMT1 or EHMT2 overexpression cells were obtained by using lentivirus to infect cells with either TRE_EHMT1_hygromycin or TRE_EHMT2_hygromycin. To produce lentivirus, 500,000 Lenti-X 293T cells were seeded on 6-well plate. Twenty-four hours later, the cells were transfected with packaging plasmids (0.5µg pCMV-VSV-G, 1.1µg pD8.9), 0.85µg transfer plasmid, 7.35µg of PEI MAX (Polysciences Inc., Cat. #24765-1) and 10mM HEPES. Media was changed 24 hours later. Supernatant with virus was collected 48 and 72 hours post transfection. The supernatant was filtered using a 0.45µm PES Filter membrane (Whatman Uniflo, Cat. #9914-2504). Cells were then infected using 1mL of supernatant with 5µg/mL polybrene (EMD Millipore, Cat. #TR-1003-G).

Media was changed 24 hours later. 48 hours post transduction, cells were selected using 200μg/ml Hygromycin B (Biosciences, Cat. #31282-04-9). Cells were selected until death of non-transduced cells. Once cells were selected, they were induced for EHMT1 or EHMT2 expression using Doxycycline (1µg/ml, TCI, Cat. #D4116) for four days and collected. Cells were immunoblotted for EHMT1 (Bethyl Laboratories, Cat. #A301-642A; dilution 1:500), EHMT2 (Cell Signaling, Cat. #3306; RRID:AB_2097647; dilution 1:1000), alpha-tubulin (Cell Signaling Cat. #3873; RRID:AB_1904178; dilution 1:3000), and H3K9me2 (Cell Signaling Cat. #4658; RRID:AB_10544405; diluted 1:1000).

### CRISPRi-mediated silencing of RIGI and MDA5

For CRISPR-mediated silencing (e.g., CRISPRi) of RIGI and MDA5, a PEO1-OR-dCas9-KRAB-MeCP2 stable line was first generated using the PiggyBac system (System Bioscience). The PiggyBac Donor plasmid, PB-CAGGS-dCas9-KRAB- MeCP2 was co-transfected with the Super PiggyBac transposase expression vector (SPBT) into PEO1-OR cells using Neon Transfection System (Thermo Fisher, Cat. #MPK5000). The pB-CAGGS- dCas9-KRAB-MeCP2 construct was a gift from Alejandro Chavez & George Church (Addgene plasmid # 110824). 24 hours post-transfection, cells were treated with Blasticidin (25μg/ml) to select for integration of the dCas9 expression cassette, and selection was maintained for 10 days. CRISPR gRNAs specific to genes of interest (i.e., 0 predicted off target sequences) were selected using pre- computed CRISPR target guides available on the UCSC Genome Browser hg38 assembly, and complementary oligos were synthesized by Integrated DNA Technologies. Complementary oligos were designed to generate BstXI and BlpI overhangs for cloning into PB-CRISPRia, a custom PiggyBac CRISPR gRNA expression plasmid based on the lentiviral construct pCRISPRia (a gift from Jonathan Weissman, RRID:Addgene 84832). Complementary gRNA-containing oligos were hybridized and phosphorylated in a single reaction, then ligated into a PB-CRISPRia expression plasmid linearized with BstXI and BlpI (New England Biolabs, Cat. #R0113L, #R0585L). Chemically competent Stable E. Coli (New England Biolabs, Cat. #C3040H) was transformed with 2 μL of each ligation reaction and resulting colonies were selected for plasmid DNA isolation using the ZymoPure Plasmid miniprep kit (Zymo Research, Cat. #D4209). Each cloned gRNA sequence-containing PB- CRISPRia plasmid was verified by Sanger sequencing (Quintara Bio). To generate CRISPRi stable lines, PB-CRISPRia gRNA plasmids were co-transfected with the PiggyBac transposase vector into the HCT116 dCas9-KRAB-MeCP2 polyclonal stable line. The following number of uniquely-mapping gRNA plasmids were designed per target based on the pre-computed UCSC hg38 CRISPR target track: GFP (1), RIGI (3), MDA5 (3) (Table S2). The same total amount of gRNA plasmid was used for transfections involving one or multiple gRNAs. 24 hours post-transfection, cells were treated with Puromycin (1*μ*g/ml) to select for integration of the sgRNA expression cassette(s). Selection was maintained for 5 days prior to transcriptional analyses.

### Compounds and Inhibitors

Olaparib was obtained from LC Laboratories (Cat. #09201), UNC0642 from MedChem Express (Cat. #HY-13980), and EZM8266 was obtained from Epizyme. For all cell lines, a dose response was performed for each compound to determine the minimal dose required for desired function and the maximum dose before acute cell defects were observed.

### RNA-sequencing

Sequencing libraries were prepared from RNA harvested from treatment or transfection replicates. Total RNA was extracted using the Quick-RNA Miniprep Plus Kit (Zymo Research, Cat. #D4209). Ribosome depletion and library preparation was performed using the Qiagen Fast-select (Cat. #334375) and KAPA BioSystems mRNA HyperPrep Kit (Cat. #KK8581) according to the manufacturer’s protocols. Briefly, 500ng of RNA was used as input, and KAPA BioSystems single-index adapters were added at a final concentration of 1.5mM. Purified, adapter-ligated library was amplified for a total of 11 cycles following the manufacturer’s protocol. The final libraries were pooled and sequenced on an Illumina NovaSeq 6000 (University of Colorado Genomics Core; RRID:SCR_021984) as 150 bp paired-end reads.

### CUT&RUN

CUT&RUN was performed using the CUT&RUN Assay Kit (Cell Signaling Technology, Cat. #86652) using the manufacturer’s protocol. 100,000 cells were used for each Input or IP sample. Input samples were sonicated using a Branson Sonifier 250. To achieve fragmentation with a peak at 300 bp, ice cold cells were pulsed 7 x 10 seconds each at 30% power, with one-minute incubations on ice between each pulse. IP was performed using 2μL EHMT2/G9A antibody (Cell Signaling Technology, Cat. #3306; RRID: AB_2097647), 2μL H3K4me3 antibody (Cell Signaling Technology, Cat. #9751; RRID: AB_2616028), or 5μL isotype control antibody (Cell Signaling Technology, Cat. #66362; RRID: AB_2924329). According to the manufacturer’s protocol, Sample Normalization Spike-In Yeast DNA was diluted 1:500 in nuclease-free water, and 5μL (10pg) of diluted DNA was added to 1X Stop Buffer prior to DNA digestion and diffusion. IP and Input DNA were purified using ChIP DNA Clean & Concentrator columns (Zymo Research, Cat. #D5205). Elution volume was 30μL. Prior to library preparation, all samples were analyzed for concentration and fragment size by the University of Colorado Pathology Shared Resource using an Agilent Tapestation with D1000 tape. Sequencing libraries were prepared using 5 ng IP DNA or 50 ng Input DNA. SimpleChIP ChIP-seq Multiplex Oligos for Illumina - Single Index Primers (Cell Signaling Technology, Cat. #29580) and the SimpleChIP ChIP-seq DNA Library Prep Kit for Illumina (Cell Signaling Technology, Cat. #56795) were used with protocol modifications for CUT&RUN as recommended by Cell Signaling. The second step of End Prep was performed for 30 minutes at 50 °C, rather than 65 °C. AMPure XP beads (Beckman Coulter, Cat. #A63880) in cleanup steps were used at a 1.1X volume, rather than 0.9X. After washing and prior to resuspension, beads were air dried for 3 minutes. To prevent large fragment amplification during PCR enrichment of adaptor-ligated DNA, the Anneal and Extension step is 65 °C for 15 seconds, rather than 75 seconds. For IP samples, 12 cycles were performed. For Input samples, 6 cycles were performed. To achieve higher final concentration, elution following PCR amplification was performed in 17μL 10 mM Tris-HCL pH 8.5, with 15 μL final collection volume. Library sequencing was performed on an Illumina NovaSEQ 6000 by the University of Colorado Genomics and Microarray Core (RRID:SCR_021984).

### CUT&TAG

CUT&TAG was performed following the protocol developed by Kaya-Okur et al^49^ with the following specifications: 500,000 (option A) PEO1-OR cells (treated with DMSO control or 2*μ*M Olaparib + 1*μ*M UNC0642 for 72 hours) were used as starting input. The following antibodies were used for primary antibody binding: H3K27Ac (EMD Millipore, Cat. #MABE647; RRID:AB_2893037), H3K4me2 (Epicypher, Cat. #13-0027), no primary antibody (negative control). Guinea Pig anti-Rabbit IgG (Antibodies, Cat. #ABIN101961; RRID:AB_10775589) was used for secondary antibody binding. For CUTAC (using H3K4me2 antibody), the CUTAC tagmentation buffer (10 mM TAPS, 5 mM MgCl2) was used instead of the CUT&TAG tagmentation buffer, and incubated at 37 degrees (step 35iv) for 20 minutes instead of 1 hour. Library sequencing was performed on an Illumina NovaSEQ 6000 by the University of Colorado Genomics and Microarray Core (RRID:SCR_021984).

### Processing of sequencing data

Reads obtained from our own datasets were reprocessed using a uniform analysis pipeline. FASTQ reads were evaluated using FastQC (v0.11.8) and MultiQC (v1.7), then trimmed using BBDuk/BBMap (v38.05). For CUT&RUN and CUT&TAG datasets, reads were aligned to the hg38 human genome using BWA (v0.7.15) and filtered for uniquely mapping reads (MAPQ > 10) with samtools (v1.10). CUT&RUN and CUT&TAG peak calls were generated using MACS2 in paired-end mode using a relaxed p-value threshold without background normalization (-- format BAMPE --pvalue 0.01 --SPMR -B --call-summits). Bedtools (v2.28.0) was used to merge peaks across the two modes of peak calling for each sample. Heatmap was generated using deepTools (v.3.0.1). RNA-seq reads were aligned to hg38 using hisat2 (v2.1.0). Bigwig tracks were generated using the bamCoverage function of deepTools (v3.0.1).

### Differential analysis using DESeq2

For RNA-seq samples, gene count tables were generated using featureCounts from the subread (v1.6.2) package with the GENCODE v34 annotation gtf to estimate counts at the gene level, over each exon (including -p to count fragments instead of reads for paired-end reads; -O to assign reads to their overlapping meta-features; -s 2 to specify reverse-strandedness; -t exon to specify the feature type; -g gene_id to specify the attribute type). To quantify TE expression at the family level, RNA-seq samples were first re-aligned to hg38 using hisat2 with -k 100 to allow multi-mapping reads and --no-softclip to disable soft-clipping of reads. TEtranscripts (v2.1.4) was then used in multi-mapping mode with the GENCODE v34 annotation gtf and hg38 GENCODE TE gtf to assign count values to both genes and TE elements. All count tables were processed with DEseq2 (v1.32.0). Normalized count values were calculated using the default DEseq2 transformation. R packages ggplot2 (v3.3.2), ggrepel (v0.8.2) and apeglm (v1.8.0) were used to visualize differentially expressed genes or TEs.

### TE colocalization analysis

To determine TE family enrichment within regulatory regions, we used GIGGLE (v0.6.3) (Layer et al. 2018) to generate a genomic interval index of all TE families in the hg38 human genome, based on Dfam v2.0 repeat annotation (n=1315 TE families). Regulatory regions (e.g., ATAC, ChIP-Seq, or CUT&RUN peaks) were queried against the TE interval index using the GIGGLE search function (-g 3209286105 -s). Results were ranked by GIGGLE enrichment score, which is a composite of the product of −log10(P value) and log2(odds ratio).

### Multispectral Immunohistochemistry (mIHC) and Image Analysis

Multispectral IHC analyses were performed using Vectra Automated Quantitative Pathology Systems (Akoya Biosciences) as described previously^50^. Tissues were formalin-fixed, paraffin embedded, and sectioned onto slides. For Fig. 5L-P, slides were sequentially stained with antibodies specific for FOXP3 (Cell Signaling, Cat. #12653S), WT1 (Novus, Cat. #110-600011), CD11c (Cell Signaling, Cat. #97585S), Ly6g (Cell Signaling, Cat. #87048S), CD3 (Cell Signaling, Cat. #D4V8L), B220 (BD Pharm, Cat. #550286), Ki67 (ThermoFisher, Cat. #RM-9106-S), F480 (Cell Signaling, Cat. #30325S), CD49b (Invitrogen, Cat. #MA5-32306), CD31 (Cell Signaling, Cat. #77699S), FOXP3 (R&D, Cat. #MAB8214), CD8 (Cell Signaling Cat. #98941S), and CD4 (Invitrogen, Cat. #14-9766-82). For Fig. 6D-E, slides were sequentially stained with antibodies specific for CD19 (Leica, Cat. #PA0843), Granzyme B (Invitrogen, Cat. #MA1-35461), CD4 (Leica, Cat. #PA0427), CD31 (Leica, Cat. #PA0414), FOXP3 (Abcam, Cat. #AB20034), CD8 (Dako/Agilent, Cat. #M703), CD68 (Dako/Agilent, Cat. #M0814), cytokeratin (Dako/Agilent, Cat. #M3515). For Fig. 7A-B, J-O, slides were sequential stained with antibodies specific for pSTAT1 (Abcam, Cat. #ab30645), MAVS (Invitrogen, Cat. #MA5-26963), IRF9 (Sigma, Cat. #HPA001862), CD3 (Leica, Cat. #PA0553), H3K9me2 (Abcam, Cat. #ab1220), cytokeratin (Dako/Agilent, Cat. #M3515), and DAPI. For Fig. 7C-I, slides were sequentially stained with antibodies specific for gH2AX (Abcam, Cat. #ab2893), cleaved caspase 3 (Cell Signaling, Cat. #9664L), PD1 (Abcam, Cat. #ab52587), PDL1 (Cell Signaling, Cat. #13684S), LAG3 (Abcam, Cat. #ab180187), Ki67 (ThermoFisher, Cat. #RM-9106-S), CD3 (Leica, Cat. #PA0553), cytokeratin (Dako/Agilent, Cat. #M3515), and DAPI. All antibody details are provided in Table S3. All slides were de-identified and imaged by the Human Immune Monitoring Shared Resource (HIMSR) core (RRID:SCR_021985) on Akoya Biosciences Vectra Polaris scanner. Regions of interest (ROIs) were selected, and multispectral images were collected with the 20x objective. A training set of 9 representative images was used to train analyses algorithms for tissue and cell segmentation and phenotyping using inForm software (Akoya Biosciences). Representative autofluorescence was measured on an unstained control slide and subtracted from study slides. Total tumor area, total cell count, and cell densities of positive and negative cells for each phenotype were graphed and compared in GraphPad Prism 9. Statistical analyses were performed using a multiple comparison One-way ANOVA test.

### Flow cytometry

Mouse peripheral blood were collected from submandibular vein. The cells were treated with red blood cell lysis buffer (0.832% NH4Cl, 0.1% NaHCO3, 0.02% EDTA) then incubated with the antibodies, CD45.2-FITC, CD3-PE, CD19-BV421, CD4-PE-Cy7 and CD8-APC (Biolegend). Stained cells were analyzed using Penteon (NovoCyte), cytometry data were analyzed using FlowJo software (TreeStar).

### Cell viability assay

Cell lines were seeded in a 96 well plate at 1000 cells per well and treated with no treatment, DMSO, 2*μ*M Olaparib, 1*μ*M UNC0642, or 2*μ*M Olaparib + 1*μ*M UNC0642. Cell media and drugs were changed every 2 days for 7 days. Cell viability of each sample in triplicates at various time points was measured using a luminescence assay of ATP (CellTiter-Glo 2.0 Assay, Promega, Cat. #G9242) following manufacturer’s protocol. Statistical analysis was done using a multiple comparison One-way ANOVA test.

### Colony formation assay

Cell lines were seeded and treated with increasing Olaparib doses as described previously in Yamamoto et al, 2019^51^. Cell medium and Olaparib were changed every 2 days for 12 days. Colonies were fixed (10% methanol/10% acetic acid) and stained with 0.4% crystal violet. Crystal violet was dissolved in fixative and absorbance was measured at 570 nm. Assays were performed in technical triplicate before reporting data.

### Reverse-transcriptase quantitative polymerase chain reaction

RNA was isolated from cells with the RNeasy Mini Kit followed by on-column DNase digest (Qiagen, Cat. #74004). mRNA expression was determined using SYBR green Luna Universal One-step RT-PCR kit (New England Biolabs, Cat #E3005L) with a BioRad CFX96 thermocycler. β-2-Microglobulin (*B2M*) and 18S rRNA were used as internal controls as stated in figured legends. All primer sequences are provided in Table S4.

### Immunoblotting

Total protein was extracted with radioimmunoprecipitation assay buffer (150 mM NaCl, 1% TritionX-100, 0.5% sodium deoxycholate, 0.1% sodium dodecyl sulfate [SDS], 50 mM Tris pH 8.0) supplemented with Complete EDTA-free protease inhibitors (Roche, Cat. #4693132001), 5mM NaF, and 1mM Na3VO4. Nuclear extraction was performed by suspending cells in a hypotonic buffer (10mM HEPES-KOH pH 7.9, 1.5mM MgCl2, 10mM KCl, 1mM DTT, 1x Halt Protease Inhibitor). After dounce homogenization and centrifugation, the resulting nuclear pellets were suspended in a hypertonic buffer (20mM HEPES-KOH pH 7.9, 25% Glycerol, 1.5mM MgCl2, 0.6M KCl, 0.2mM EDTA, 1mM DTT, 1x Halt Protease Inhibitor). Protein was separated on an SDS polyacrylamide gel electrophoresis and transferred to polyvinylidene fluoride membrane. Primary antibody incubation was performed overnight at 4C. Secondary goat anti-rabbit (IRDye 680RD or IRDye 800CW, LI-COR, Cat. #92568071; RRID: AB_2721181 or Cat. #926-32211; RRID: AB_621842; 1:20,000) and goat anti-mouse (IRDye 680RD or IRDye 800CW, LI-COR, Cat. # 926-68070; RRID: AB_10956588 or Cat# 925-32210; RRID: AB_2687825; 1:20,000) antibodies were applied for 1 hour at room temperature. Blots were visualized using the Licor Odyssey Imaging System and ImageStudio software (V4).

### Immunofluorescence

Cells were seeded on pretreated glass coverslips. 24 hours later plates were fixed in 4% paraformaldehyde. Cells were permeabilized in 0.2% TritonX in PBS and incubated in primary antibody for two hours at room temperature. Cells were rinsed with 1% TritonX and incubated in secondary antibody for one hour at room temperature. Glass slides are mounted with SlowFade Dapi and sealed with nail polish. Images were taken using a Nikon DS-Ri2 and at least 200 cells were quantified.

### In vivo studies

All mouse experiments were approved by the Institutional Animal Care and Use Committee (IACUC protocol No. 569). For all studies six- to eight-week-old C57BL/6J mice were purchased from Jackson Laboratories (strain #000664), and these mice were intraperitoneally (i.p.) injected with 5 × 10^6^ Olaparib-resistant ID8 p53^-/-^ BRCA2^-/-^ cells. Tumor cells were tagged with GFP/luciferase to facilitate tracking via In Vivo Imaging Software (IVIS, Perkin Elmer)^51, 52^. IVIS imaging conduted at Small Animal Imaging Shared Resource (RRID:SCR_021980). Mice were IVIS scanned weekly starting 7 days after ID8 cell injection (day 0), tumor burden is reported as total flux (photons/second). For all studies, mice were randomized into treatment groups based on day 0 IVIS total flux, and the 28-day treatment regimens began on day 1. Mice were euthanized and necropsied on day 29 in all experiments. EZM8266 was made daily in 0.1% Tween80 (Sigma-Aldrich, Cat. no. P1754), 0.5% methylcellulose (Sigma-Aldrich, Cat. no. 64632) in sterile water as a vehicle. Mice were weighed twice per week, and bodyweight served as a surrogate for drug toxicity.

*Experiment 1(Figure 5A)*: Mice were randomized into four treatment groups by day 0 IVIS total flux: Vehicle control (n = 10), 50 mg/kg Olaparib (LC Laboratories, Cat. #09201, n = 10), 300 mg/kg EZM8266 (n = 9), or Olaparib + EZM8266 (n = 10). All treatments were administered daily via oral gavage. Immediately prior to death, we collected ∼100 μl blood via submandibular puncture (n = 5 Vehicle control, n = 5 Olaparib, n = 5 EZM8266, and n = 4 Olaparib + EZM8266) and stored the samples in lithium-heparin coated microtubes (Sarstedt AG % Co. KG, Cat. #41.1393.105) at 4°C. Mouse Complete Blood Count (CBC) was performed on lithium heparin anti-coagulated blood samples using an automated analyzer (HemaTrue, Heska Corp, Loveland, CO) in the Comparative Pathology Shared Resource at University of Colorado Anschutz Medical Campus.

*Experiment 2 (Figure S4)*: Mice were randomized into four treatment groups by day 0 total flux: Vehicle control (n = 5), 75 mg/kg EZM8266 (n = 5), 150 mg/kg EZM8266 (n = 5), or 300 mg/kg EZM8266 (n = 5). Vehicle or EZM8266 was administered daily via oral gavage for 28 days.

*Experiment 3 (Figure 6F)*: On days -3, -2, and -1, mice received a daily i.p. injection of either 500 μg anti-CD8a antibody (BioXCell, Cat. no. BE0004-1; RRID:AB_1107671) or 500 μg *InVivo*MAb rat IgG2a isotype control, anti-trinitrophenol (BioXCell, Cat. #BE0079). *InVivo*Pure pH 6.5 dilution buffer (BioXCell, Cat. #IP0065) was used as a vehicle for both antibodies. On day 0, CD8 T cell depletion was validated via flow cytometry (Figure S5A-B), and mice within each antibody group were randomized into four treatment groups based on IVIS total flux: IgG2a + Vehicle control (n = 9), Anti- CD8a antibody (n = 10), 150 mg/kg EZM8266 (n = 9), and Anti-CD8a + EZM8266 (n = 10). EZM8266 was administered daily via oral gavage, and 250 μg anti-CD8a antibody was administered via i.p. injection twice per week starting on day 1.

### Data availability

All RNA-seq sequencing and epigenomic profiling data are deposited in NCBI GEO Accession: GSE224062 and GSE225338.

## ACKNOWLEDGMENTS

We acknowledge philanthropic contributions from Kay L. Dunton Endowed Memorial Professorship In Ovarian Cancer Research, the McClintock-Addlesperger Family, Karen M. Jennison, Don and Arlene Mohler Johnson Family, Michael Intagliata, Duane and Denise Suess, Mary Normandin, and Donald Engelstad. Chuong E is supported by the Alfred P. Sloan Foundation, the David and Lucile Packard Foundation, and the Boettcher foundation. Bitler B is supported by The Department of Defense (Bitler, OC170228, OC200302, OC200225), The American Cancer Society (Bitler, RSG-19-129-01- DDC), The Ovarian Cancer Research Alliance (Bitler B) and National Institutes of Health (Bitler B, R37CA261987; Arnoult N R01CA266100; Chuong E R35GM128822). Arnoult N. was supported by the NIH (R35GM143108), the Boettcher Foundation, the V Foundation for Cancer Research, and the Glenn Foundation and American Federation for Aging Research. Watson ZL is supported by the National Institutes of Health (R03CA249571) and the Department of Defense Ovarian Cancer Research Program (OC210257). The University of Colorado Cancer Center Support Grant (P30CA046934). We acknowledge support from the University of Colorado Cancer Center and the MCDB. We acknowledge Etienne Danis for his contribution to the CUT&RUN analysis. Thank you to Hei-yong Grant Lo for his unwavering support.

---

**Figure S1:**
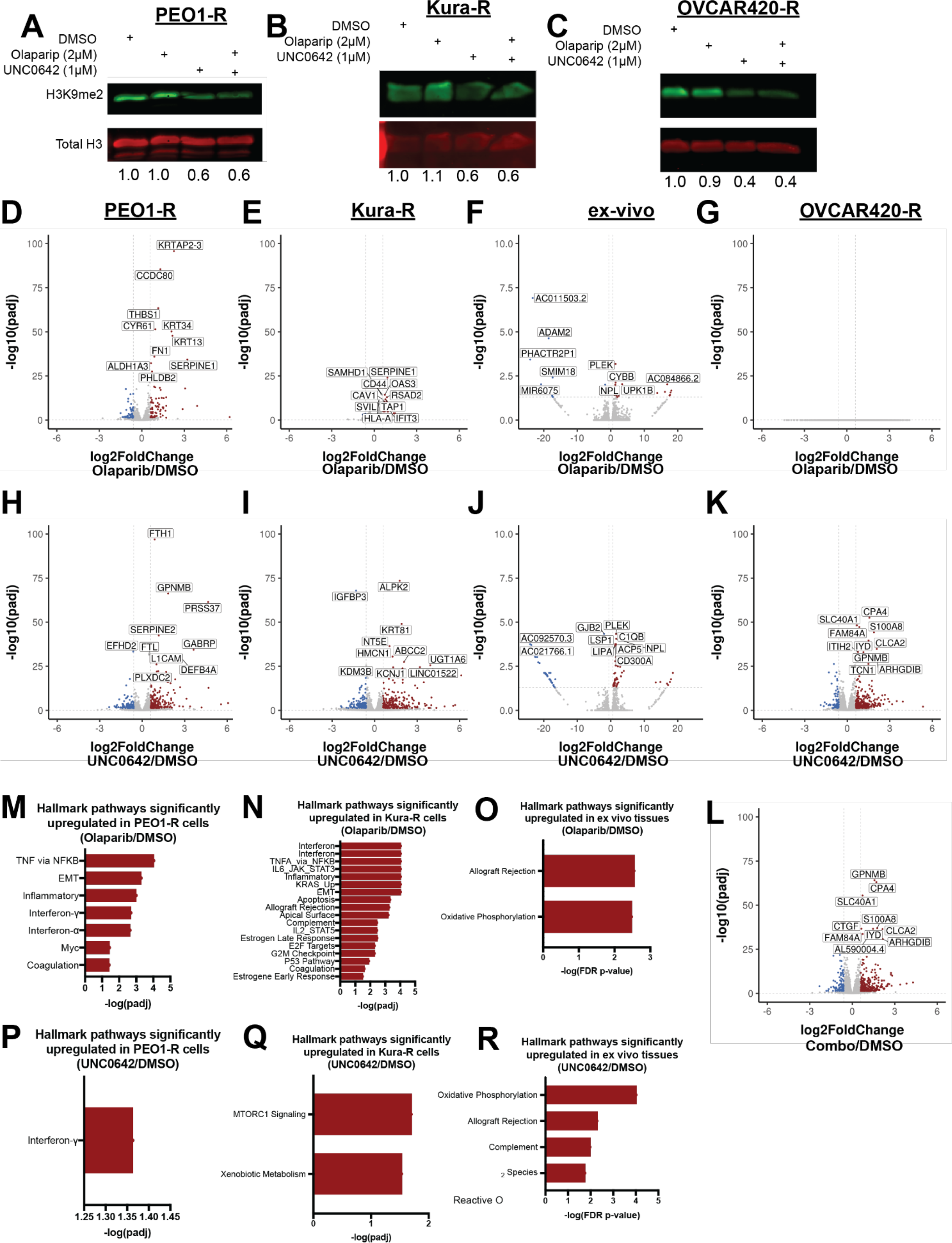
Combinatory EHMT and PARP inhibition induces transcriptional reprogramming and upregulation of interferon pathways. **A-C)** Protein from PEO1-R (A), Kura-R (B), and OVCAR420-R (C) treated with DMSO, Olaparib, UNC0642, or Olaparib/UNC0642 used for immunoblot against H3K9me2. Loading control, total Histone 3. Quantification, below. **D-G)** Volcano plot of differentially regulated genes in PEO1-R (D), Kura-R (E), patient-derived *ex vivo* tissues (F), and OVCAR420-R (G) cells treated with Olaparib compared to DMSO control. **H-K)** Volcano plot of differentially regulated genes in PEO1-R (H), Kura-R (I), patient-derived *ex vivo* tissues (J), and OVCAR420-R (K) cells treated with UNC0642 compared to DMSO. **L)** Volcano plot of differentially regulated genes in OVCAR420-R (L) cells treated with Olaparib/UNC0642 compared to DMSO. **M-O)** List of hallmark pathways upregulated in PEO1-R (M), Kura-R (N), and patient-derived *ex vivo* tissues (O) treated with Olaparib compared to DMSO. **P-R)** List of hallmark pathways upregulated in PEO1-R (P), Kura-R (Q), and patient-derived *ex vivo* tissues (R) treated with UNC0642 compared to DMSO.0.8

**Figure S2:**
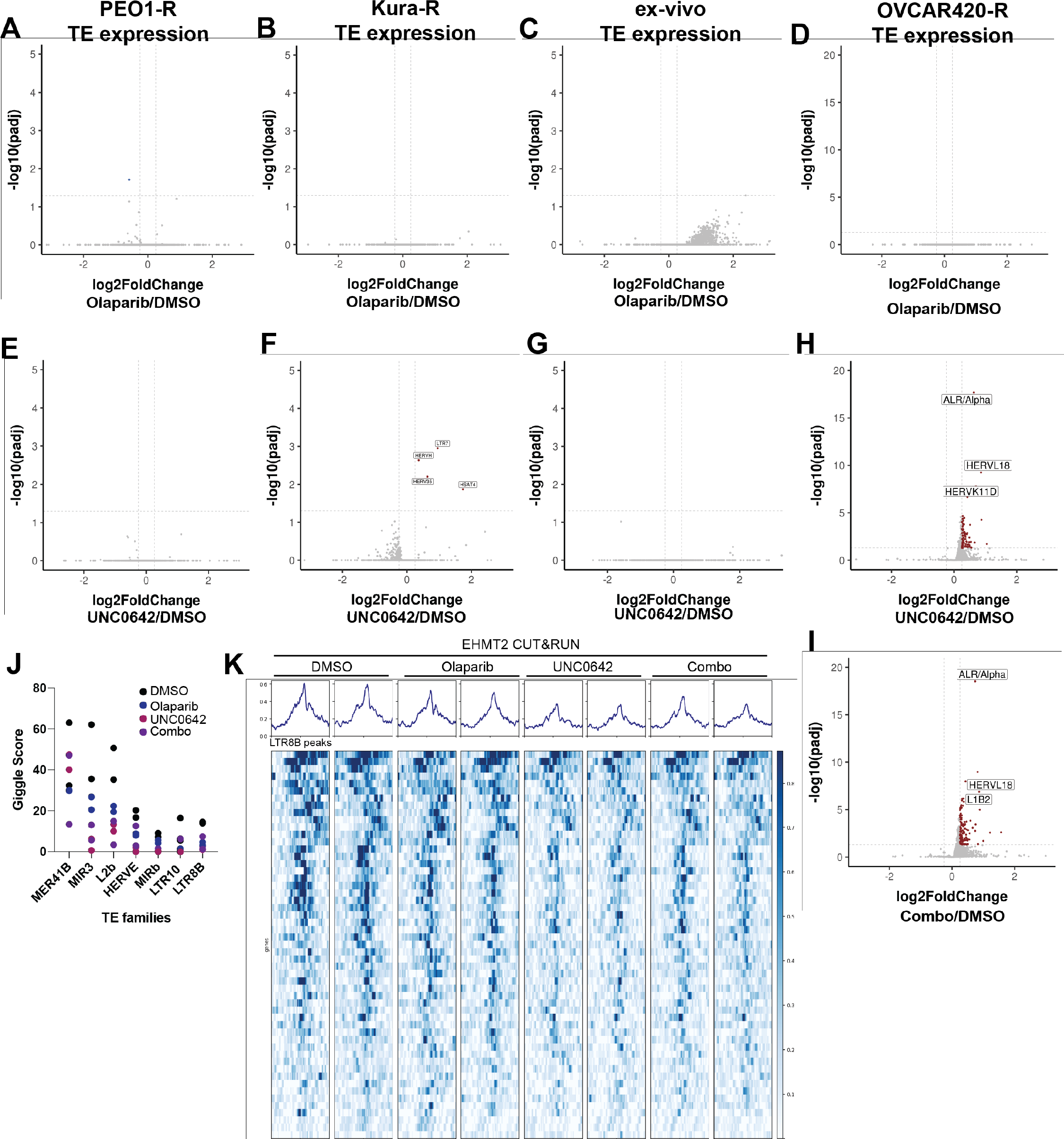
Combination therapy induces expression of transposable elements. **A-D)** Volcano plots of TE families with differentially expressed transcripts in PEO1-R (A), Kura-R (B), patient-derived *ex vivo* tissues (C), and OVCAR420-R (D) cells treated with Olaparib compared to DMSO. **E-H)** Volcano plots of TE families with differentially expressed transcripts in PEO1-R (E), Kura-R (F), patient-derived *ex vivo* tissues (G), and OVCAR420-R (H) cells treated with UNC0642 compared to DMSO. **I)** Volcano plots of TE families with differentially expressed transcripts in OVCAR420-R cells treated with Olaparib/UNC0642 compared to DMSO. **J)** Giggle score of TE families found to be enriched in EHMT2 CUT&RUN experiment. **K)** LTR8B signals at EHMT2-bound loci.

**Figure S3.**
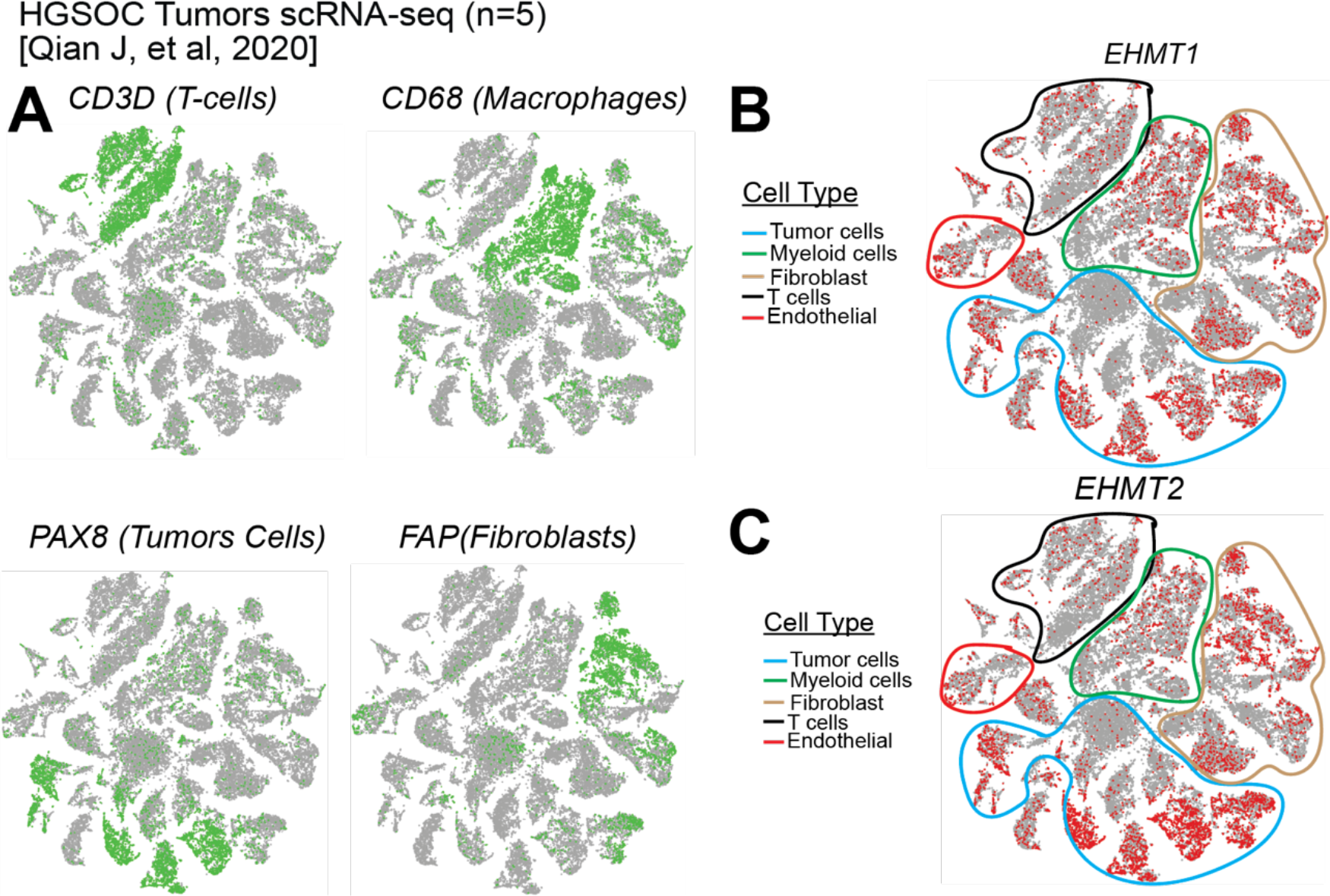
*EHMT1* and *EHMT2* expression in HGSOC tumors. scRNA-sequencing from five independent HGSOC tumors clustered based on transcriptional profiles. **A)** Expression of cell type specific markers. Expression of **B)** EHMT1 and **C)** EHMT2 in the different cell type clusters.

**Figure S4.**
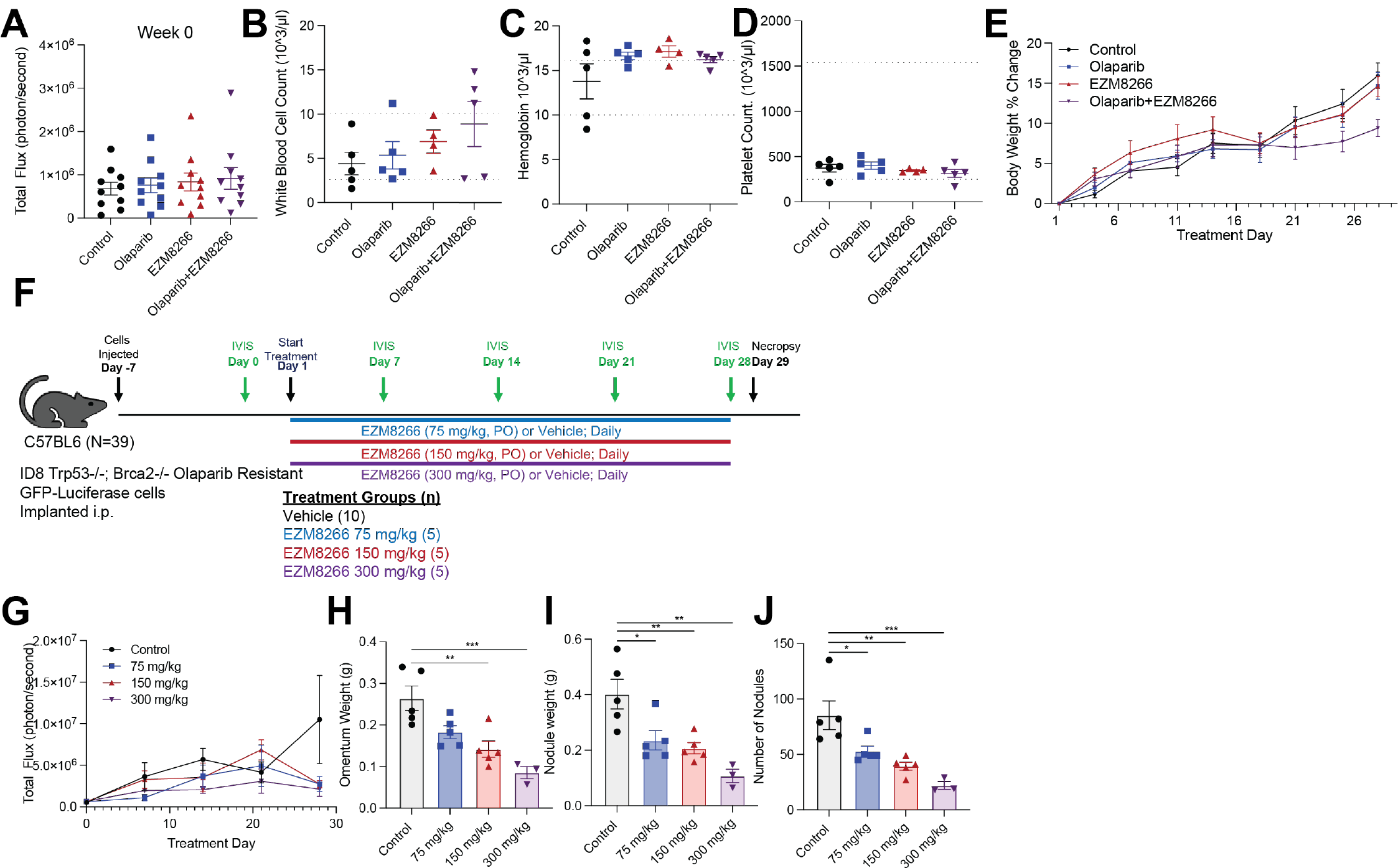
EHMT inhibition is well-tolerated and the anti-tumor response is dose-dependent. **A)** Randomization of animals based on total flux at beginning of the study (Study design shown in Fig. 5A). **B)** White blood cell count of treated animals. **C)** Hemoglobin count of treated animals. **D)** Platelet count of treated animals. **E)** Body weight over the course of treatment. **F)** Tumor response is dose dependent. Study design and population size indicated next to treatment (n=5/group). **G)** Tumor progression monitored via total flux. **H)** Omental weight. **I)** Solid tumor weight. **J)** Number of tumor dissemination sites. Error bars, SEM. Statistical test, multicomparison ANOVA with Tukey correction.

**Figure S5.**
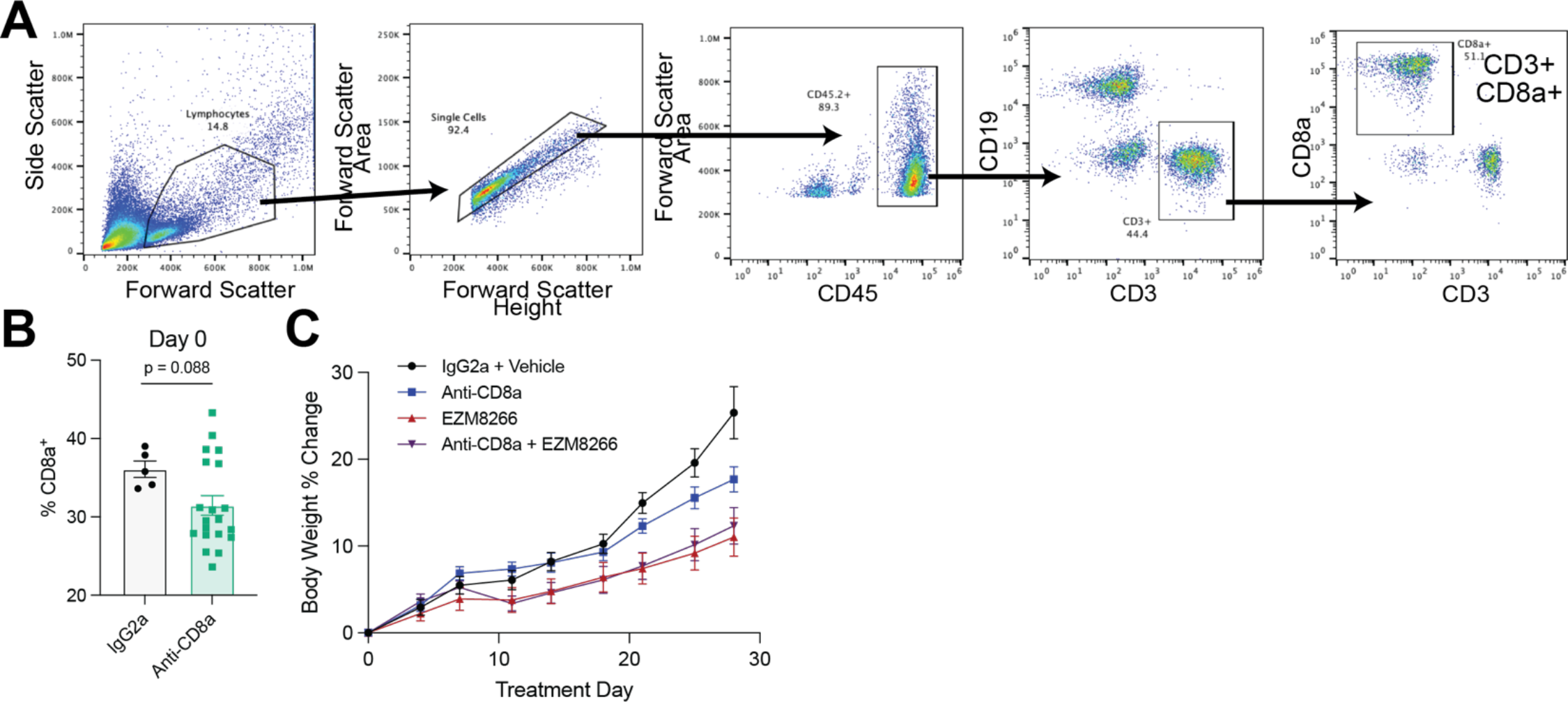
**A)** Gating strategy for CD8 depletion *in vivo* study. **B)** Serum associated T cells at the time of treatment initiation. **C)** Body weight of tumor-bearing mice over the course of treatment.

**Figure S6:**
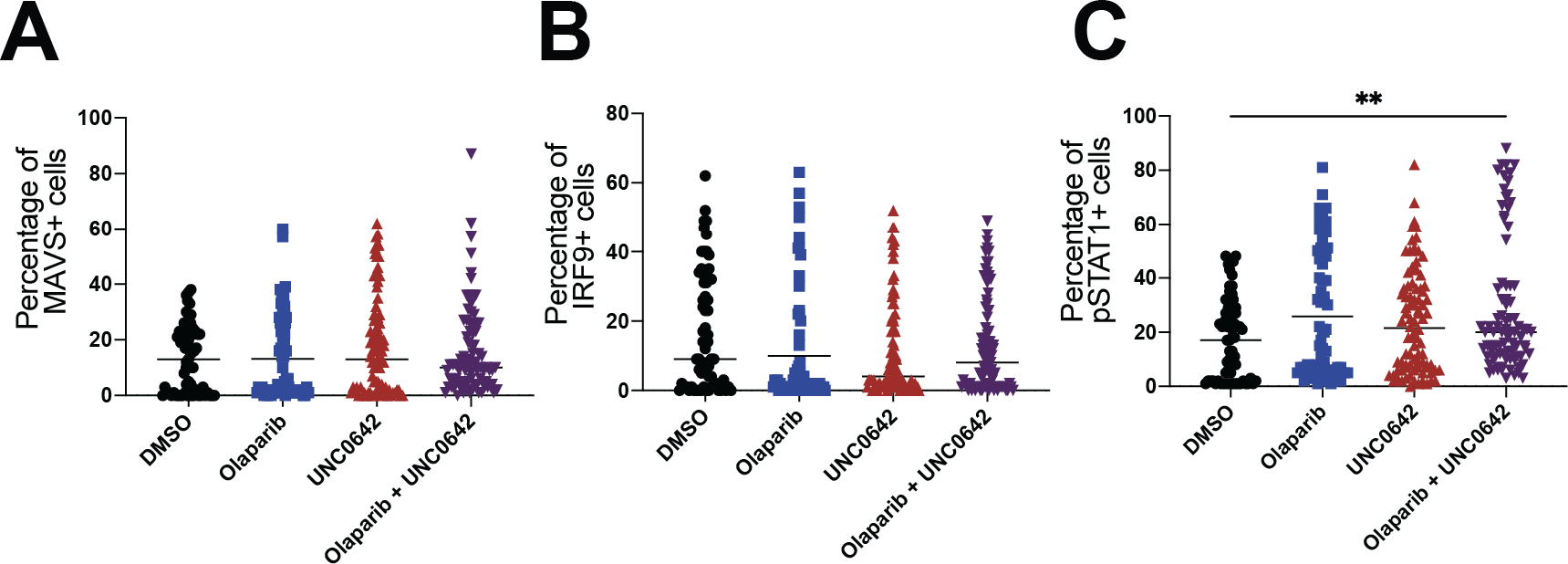
Combination therapy may contribute to T cell exhaustion and dsRNA-induced interferon response *ex vivo*. Primary ovarian cancer tumors treated with EHMT inhibitor (UNC0462) and/or Olaparib. Tumor sections were fixed and analyzed via multispectral IHC. Quantification of **A)** MAVS, **B)** IRF9, and **C)** pSTAT1. Error bars, SEM. Statistical test, multi-comparison ANOVA with Tukey correction. **p<0.01

**Supplemental Table 1:** Characterization of patient-derived ex vivo tissues obtained from the Gynecology Tissue and Fluid Bank at the University of Colorado Anschutz Medical Campus.

| GTFB Number | Diagnosis |
| --- | --- |
| 1229 | stage IIIA grade 2 endometrioid endometrial adenocarcinoma |
| 1230 | stage IIIC high grade serous fallopian tube carcinoma |
| 1236 | stage IIIB high grade serous ovarian carcinoma |

**Supplemental Table 2:**
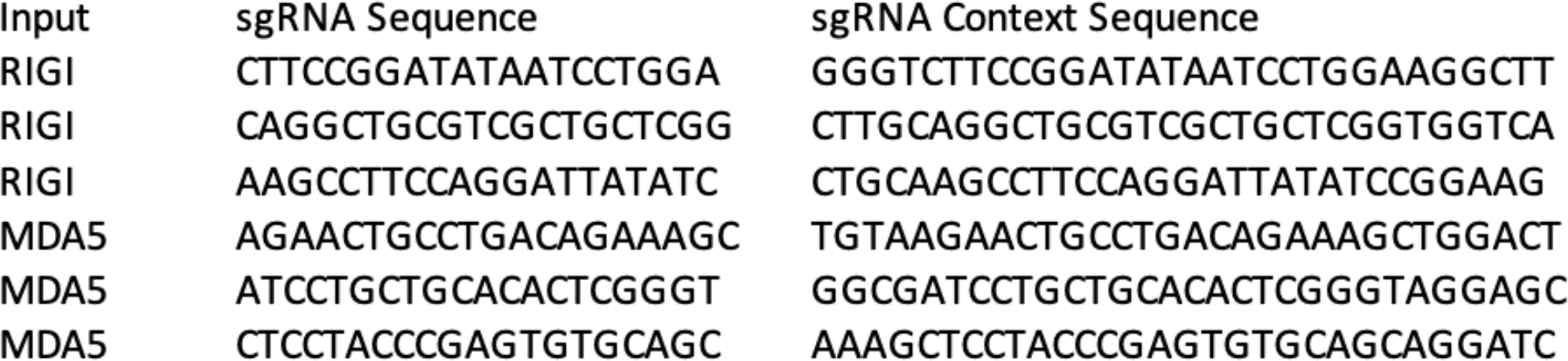
sgRNA sequences for CRISPRi-mediated silencing.

**Supplemental Table 3:**
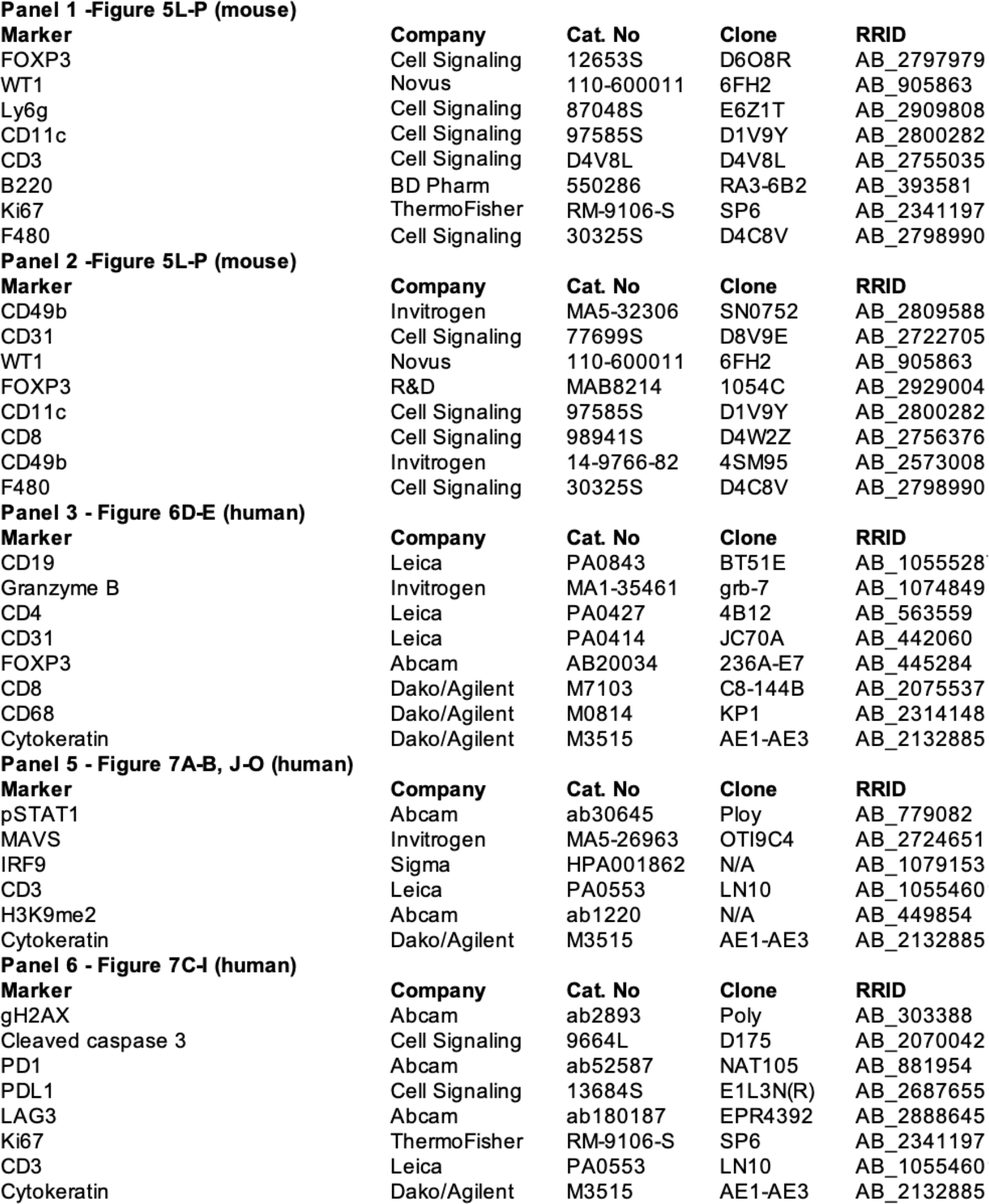
Information of antibodies used for multispectral IHC experiments

**Panel 1 -Figure 5L-P (mouse)**
| Marker | Company | Cat. No | Clone | RRID |
| --- | --- | --- | --- | --- |
| FOXP3 | Cell Signaling | 12653S | D6O8R | AB_2797979 |
| WT1 | Novus | 110-600011 | 6FH2 | AB_905863 |
| Ly6g | Cell Signaling | 87048S | E6Z1T | AB_2909808 |
| CD11c | Cell Signaling | 97585S | D1V9Y | AB_2800282 |
| CD3 | Cell Signaling | D4V8L | D4V8L | AB_2755035 |
| B220 | BD Pharm | 550286 | RA3-6B2 | AB_393581 |
| Ki67 | ThermoFisher | RM-9106-S | SP6 | AB_2341197 |
| F480 | Cell Signaling | 30325S | D4C8V | AB_2798990 |

**Panel 2 -Figure 5L-P (mouse)**
| Marker | Company | Cat. No | Clone | RRID |
| --- | --- | --- | --- | --- |
| CD49b | Invitrogen | MA5-32306 | SN0752 | AB_2809588 |
| CD31 | Cell Signaling | 77699S | D8V9E | AB_2722705 |
| WT1 | Novus | 110-600011 | 6FH2 | AB_905863 |
| FOXP3 | R&D | MAB8214 | 1054C | AB_2929004 |
| CD11c | Cell Signaling | 97585S | D1V9Y | AB_2800282 |
| CD8 | Cell Signaling | 98941S | D4W2Z | AB_2756376 |
| CD49b | Invitrogen | 14-9766-82 | 4SM95 | AB_2573008 |
| F480 | Cell Signaling | 30325S | D4C8V | AB_2798990 |

**Panel 3 - Figure 6D-E (human)**
| Marker | Company | Cat. No | Clone | RRID |
| --- | --- | --- | --- | --- |
| CD19 | Leica | PA0843 | BT51E | AB_1055528 |
| Granzyme B | Invitrogen | MA1-35461 | grb-7 | AB_1074849 |
| CD4 | Leica | PA0427 | 4B12 | AB_563559 |
| CD31 | Leica | PA0414 | JC70A | AB_442060 |
| FOXP3 | Abcam | AB20034 | 236A-E7 | AB_445284 |
| CD8 | Dako/Agilent | M7103 | C8-144B | AB_2075537 |
| CD68 | Dako/Agilent | M0814 | KP1 | AB_2314148 |
| Cytokeratin | Dako/Agilent | M3515 | AE1-AE3 | AB_2132885 |

**Panel 5 - Figure 7A-B, J-O (human)**
| Marker | Company | Cat. No | Clone | RRID |
| --- | --- | --- | --- | --- |
| pSTAT1 | Abcam | ab30645 | Ploy | AB_779082 |
| MAVS | Invitrogen | MA5-26963 | OTI9C4 | AB_2724651 |
| IRF9 | Sigma | HPA001862 | N/A | AB_1079153 |
| CD3 | Leica | PA0553 | LN10 | AB_1055460 |
| H3K9me2 | Abcam | ab1220 | N/A | AB_449854 |
| Cytokeratin | Dako/Agilent | M3515 | AE1-AE3 | AB_2132885 |

| Marker | Company | Cat. No | Clone | RRID |
| --- | --- | --- | --- | --- |
| gH2AX | Abcam | ab2893 | Poly | AB_303388 |
| Cleaved caspase 3 | Cell Signaling | 9664L | D175 | AB_2070042 |
| PD1 | Abcam | ab52587 | NAT105 | AB_881954 |
| PDL1 | Cell Signaling | 13684S | E1L3N(R) | AB_2687655 |
| LAG3 | Abcam | ab180187 | EPR4392 | AB_2888645 |
| Ki67 | ThermoFisher | RM-9106-S | SP6 | AB_2341197 |
| CD3 | Leica | PA0553 | LN10 | AB_1055460 |
| Cytokeratin | Dako/Agilent | M3515 | AE1-AE3 | AB_2132885 |

**Supplemental Table 4:**
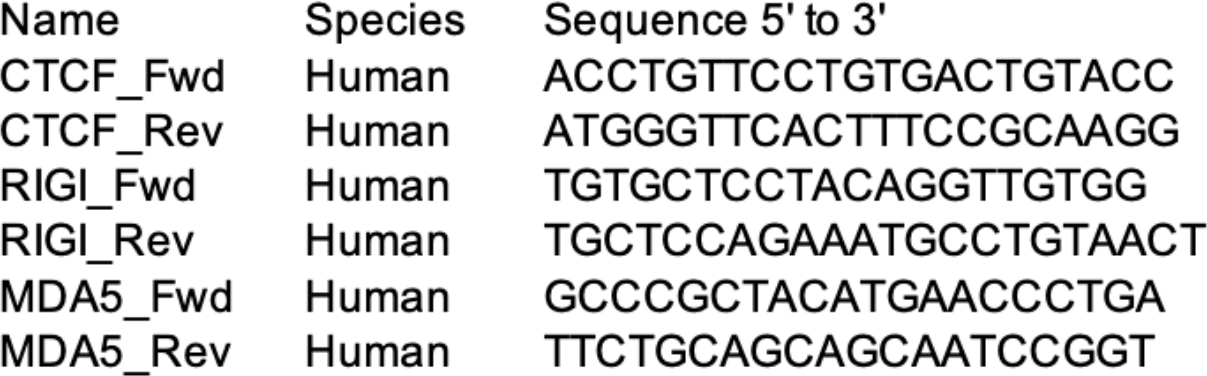
Sequences of qRT-PCR primers.

